# A telencephalon cell type atlas for goldfish reveals diversity in the evolution of spatial structure and cell types

**DOI:** 10.1101/2023.06.19.545605

**Authors:** Muhammad Tibi, Stav Biton, Hannah Hochgerner, Zhige Lin, Shachar Givon, Osnat Ophir, Tal Shay, Thomas Mueller, Ronen Segev, Amit Zeisel

**Author notes:** Correspondence to A.Z. and R.S. M.T. and S.B. contributed equally.

## Abstract

Teleost fish form the largest group of vertebrates, making them critically important for the study on the mechanisms of brain evolution. In fact, teleosts show a tremendous variety of adaptive behaviors similar to birds and mammals, however, the neural basis mediating these behaviors remains elusive. We performed a systematic comparative survey of the goldfish telencephalon; the seat of plastic behavior, learning and memory in vertebrates. We delineated and mapped goldfish telencephalon cell types using single-cell RNA-seq and spatial transcriptomics, resulting in *de novo* molecular neuroanatomy parcellation. Glial cells were highly conserved across 450 million years of evolution separating mouse and goldfish, while neurons showed diversity and modularity in gene expression. Specifically, somatostatin (SST) interneurons, famously interspersed in the mammalian isocortex for local inhibitory input, were curiously aggregated in a single goldfish telencephalon nucleus, but molecularly conserved. Cerebral nuclei including the striatum, a hub for motivated behavior in amniotes, had molecularly and spatially conserved goldfish homologues. We further suggest different elements of a hippocampal

formation across the goldfish pallium. Together, our atlas provides new insights to organization and evolution of vertebrate forebrains and may serve as a resource for the functional study underlying cognition in teleost fish.

**Teaser:** Detailed mapping of goldfish forebrain cells unwraps how 450 million years of evolution may have impacted brain function

## Introduction

The vertebrates’ telencephalon, part of the forebrain, is responsible for higher cognitive functions such as learning and memory ^1^. After massive lesion to the telencephalon, most non-mammalian vertebrates retain the capacity to perform movements, but display major deficits in predicting the outcome of future actions ^2–5^. And while other parts of the brain are conserved among vertebrates, telencephalon size and complexity vary considerably across teleost ray-finned fish, amphibians and amniotes such as reptiles, birds and mammals ^6, 7^. Teleost fish are the largest vertebrate family, represented by around 26,000 species. They diverged from the amphibian and amniotes branches of vertebrates 450 *mya.* Their study therefore provides a unique viewpoint on principles of brain evolution and structure.

However, understanding the evolution of brain structure and cell types across the vertebrates’ phylogenetic tree faces a long-debated problem in comparative neurobiology: the evolution of an outwardly-grown (everted) telencephalon, which is specific for ray-finned fish (*Actinopterygii*). The everted telencephalon, in fact, does not anatomically resemble inwardly grown (evaginated) ones of non-actinopterygian vertebrates (e.g., reptiles and mammals)^8^. This makes anatomical comparisons to well-known forebrains of tetrapod challenging. Moreover, teleost fish, like goldfish, represent the most modern group of ray-finned fish and their telencephala show great diversity in terms of brain and behavioral complexity, and adaptations to spatial ecologies and lifestyles.

This long debated problem led to a controversy about the actual brain divisions of different teleost species and their exact homologies with other vertebrates ^9, 10^. Today, there is no accepted complete mapping between several major brain regions in the teleost and amniotes. For example, the prime region for spatial cognition in amniotes is the hippocampal formation, which forms a distinct structure, where place cells are recorded ^11, 12^. According to the evagination-eversion process, the teleost counterpart is expected in the lateral pallium. Lesion studies and electrophysiological recordings found evidence for the location xof space representing cells in the lateral pallium ^5^ ^13^, but also the medial pallium ^14^ and central telencephalon ^15^.

It remains an open question to what extent brain regions in teleosts are truly homologous to comparable regions in mammals and other tetrapods.

Here, we argue that debates about the evolutionary interpretations of pallial and subpallial territories should be informed by systematic molecular evidence: Understanding the organization of the teleostean telencephalon could be greatly aided by uncovering cellular homologs. Therefore, we used an unbiased, transcriptome-wide gene expression analysis approach: We combined two transcriptome-wide technologies, scRNA-seq ^16^ and spatial transcriptomics ^17^, to generate a molecularly and spatially resolved cell type atlas for the goldfish telencephalon. We identified forebrain territories and revealed genes with axial patterned expression. In systematic comparison to the most thoroughly investigated forebrain today, the mouse’, we identified several molecularly conserved cell types and brain structures in goldfish forebrain. Finally, we also established an accessible web resource for free exploration of gene expression across cell types, and spatial context.

## Results

### Goldfish telencephalon cell type atlas

Single-cell RNA-sequencing (scRNA-seq) enables the comparison of molecular similarity and differences of cell types of a given tissue across species, purely based on genome-wide gene expression profiles (transcriptomes). Furthermore, spatial transcriptomics approaches can retain positional information of mRNA molecules across the tissue and enable the spatial mapping of gene profiles. Combining these two techniques, we generated a spatially annotated cell type atlas for the goldfish telencephalon (Fig. 1A).

**Figure 1:**
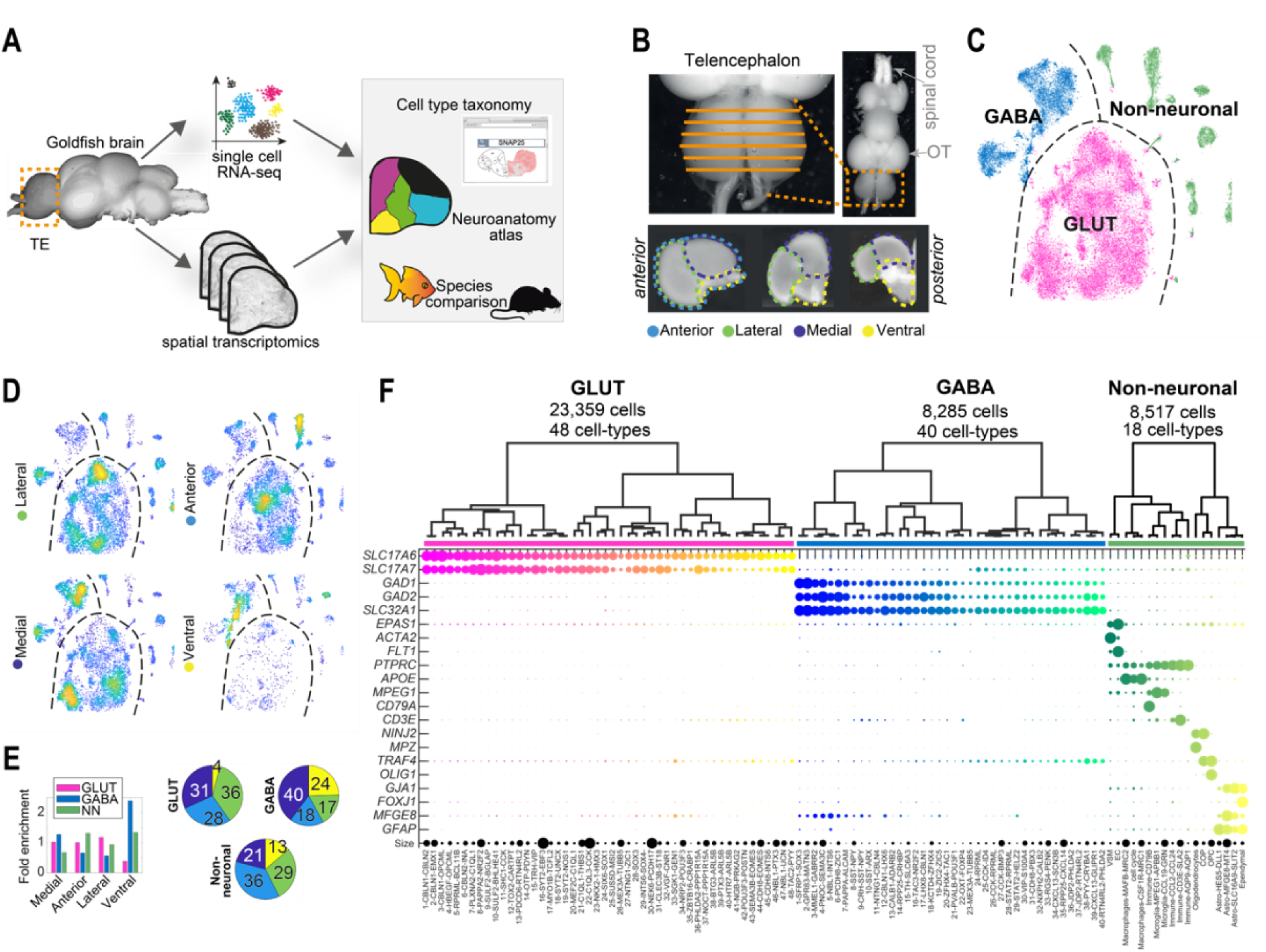
Composition of goldfish Telencephalon cells. **A** Outline of this study. Adult goldfish telencephala were analyzed in parallel using spatial- and single cell transcriptomics, integrated to a spatially mapped cell type atlas, and followed by species comparison. **B** Images of fresh dissected goldfish brain and coronal telencephalon sections, describing the microdissection scheme. **C** t-SNE visualization of all cells in the goldfish forebrain. Each dot represents a cell, colored by class. **D** Density scatter visualization on t-SNE of all cells (as in C), per microdissection origin; yellow, dense; blue, sparse. **E** Left: Contribution of four microdissections to each class; visualized as pie charts. Right: Contribution of cell class to each dissection; visualized as bar plot. **F** Dendrogram overview of all cell types; and below, top marker gene expression visualized as Dotplot. Dot size represents the percentage of cells in a cluster expressing each gene; dot color represents cell type.

First, we performed scRNA-seq on the full goldfish telencephalon (see Methods). From fresh coronal sections of total ten goldfish, we microdissected lateral, medial, ventral or anterior telencephalon (Fig. 1B). Each region was dissociated to highly viable cell suspensions, single cells encapsulated to microfluidic droplets (10x Genomics), sequenced, and mapped to the reference genome (see Methods). By their expression profile, we then assigned cells to putative cell types, summarized in Fig. 1 (see Methods).

At first, based on known markers (e.g. neurotransmitter transporters *SLC17A7*, *SLC17A6* and *SLC32A1*), we divided the ∼40,000 cells into three main classes: Glutamatergic neurons (∼23,400 cells), GABAergic neurons (∼8,300 cells) and non-neuronal cells (∼8,500 cells), (Fig. 1C). As expected, the dissection compartments (lateral, ventral, medial, anterior) differed in terms of cell type composition (Fig. 1D-E); for example, GABAergic cells dominated the ventral telencephalon. The general cell classes contained 106 defined transcriptomic cell types (Fig. 1F), where 40 were GABAergic, 48 glutamatergic, and 18 cell types were non-neuronal (i.e., glial and immune cells; Fig. S1, see also species comparison Fig. 6). We discuss neuronal cell types by a number handle according to the hierarchical clustering (i.e., **GABA1-40**, and **GLUT1-48**), additionally highlight 1-3 marker genes in the cell type names in figures (e.g., **GABA6**-PCDH8-ZIC1).

### Spatial transcriptomics allowed unbiased delineation of goldfish telencephalon territories

Single-cell RNA-seq revealed molecular expression profiles of cells within the dissected goldfish telencephalon regions, to generate the cell atlas. However, tissue dissociation for scRNA-seq came at the loss of more refined spatial information, critical for defining topological identity and possible homologies to other species. Therefore, we next performed spatial transcriptomics along the telencephalon’s anterior-posterior axis (Fig. 1A). Spatial transcriptomics (Visium, 10xGenomics) is a sequencing-based, transcriptome-wide *in situ* mRNA-detection method; at the loss of cellular resolution: The method retained positional information of mRNA molecules, per 55μm-diameter capture spots, that are arranged in an array with centers (X-Y) 100μm apart. We sampled eight 10μm coronal sections, spaced on average 100μm apart (Z), from two goldfish telencephala, each. Thus, at a pixel size of ∼100μm, the spatial transcriptomics data includes a complete map (image) of each gene’s expression, in the goldfish telencephalon.

To test if the transcriptome can reveal spatial patterns and allow us to delineate territories in the goldfish telencephalon, we performed unbiased clustering of the 6,710 Visium capture spots we sampled. We obtained 17 “spatial” clusters based on distinct molecular profiles and remapped the cluster assignment to tissue sections (see Methods and Fig. S2). Although the clustering procedure was dependent solely on gene expression, and based on a “bulk” of cells present per 55μm capture spot, we discovered clear spatial segmentation in each section. For example, in anterior sections, superficial spots were distinct from all deeper spots. This was likely due to a strong signature of *MFGE8+* astrocytes present at the pial surface (Fig. S1). More posterior, axial patterning appeared; and some smaller, spatially refined molecular territories became evident; resembling anatomical nuclei.

### Regional distribution of goldfish neurons allow telencephalon parcellation

Noting how spatial transcriptomics revealed coarse anatomical parcellations confirmed that transcriptome encodes anatomical information in the goldfish telenecephalon. And critically, in combination with the scRNA-seq data, the spatial dataset next allowed us to localize molecularly defined neuronal cell types in the telencephalon: To map the probable location of each scRNA-seq cell type to anatomical positions, we applied a custom-made algorithm on the spatial dataset. For each molecular cell type, we scored and visualized the detected expression of its top enriched genes per spot, in anterior-posterior sections of two replicate brains (Fig. S3). Since each spot of the spatial transcriptomics array potentially contains on the order of 10 cells, several cell types can map to the same spot. Therefore, we next computed the weighted sum of cell-type scores per spot and visualized the integrated mapping of 88 cell types using the cell types colormap of Fig. 1F (Fig. 2A, Fig. S4).

**Figure 2:**
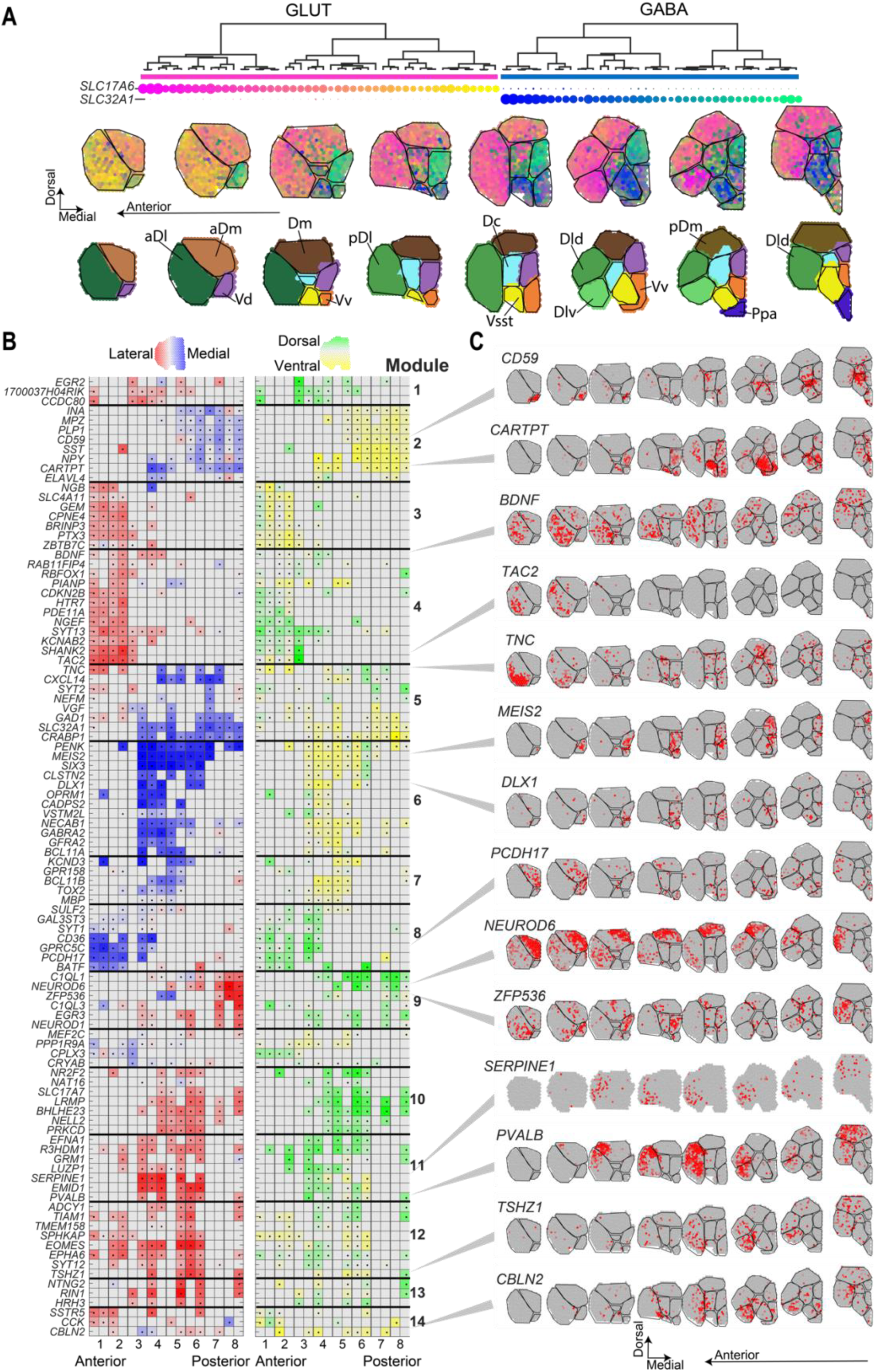
Axial and regional parcellation of goldfish telencephalon. **A** Molecular cell type-based neuroanatomy of the goldfish telencephalon. ***Top***: color scheme shown above for GABAergic and glutamatergic cell types (dendrogram); ***Center***: Eight coronal goldfish sections (goldfish 1), sampled for Visium spatial transcriptomics, overlaid with a weighted colormap that integrates all goldfish telencephalon cell types. ***Bottom***: Regional parcellation based on colormap differences above, each color indicates a different region, and suggested nomenclature annotated by similarity to Northcutt region names. **B** Heatmaps of normalized standard deviation scores for top axial pattern genes in the dorso-ventral (left) and medio-lateral (right) axes, for goldfish 1 and 2; dots indicate spatial enrichment according to axial color scheme shown above; grey without dot, no enrichment. **C** Expression of 14 axial-patterned genes across the goldfish telencephalon; grey, low; red, high.

This mapping provides a visual intuition of the dominant cell types in each spot, and provided several key insights: (1) Molecularly related cell types showed overlapping spatial patterns, (2) all territories were described by a multitude of cell types and (3) some territories were clearly distinct from neighbors with sharp borders, contrasting others with more diffuse borders.

For example, glutamatergic types **GLUT41-48** made up a closely related branch in gene expression, and all localized laterally, in a territory of the anterior telencephalon only (Fig. 2A, yellow). They showed some overlap with the neighboring branch of cell types (**GLUT33-40**); however the latter continued more dorsal, and into posterior sections. Consequently, the border between the anterior-lateral, and the more dorsal territory appeared diffused compared for example, to the sharp border defined by GABAergic clusters **24-40** (green).

Taking advantage of visual color borders, we manually drew regional outlines within each telencephalon Visium section. This resulted in a neuroanatomical delineation based on molecularly defined neuronal cell types. In most cases, we adopted regional names for these defined territories following commonly used terminology for goldfish neuroanatomy ^18, 19^ (Fig. 2A, bottom). In fact, our anatomical parcellation was astoundingly consistent with traditionally recognized delineations that were mainly based on cytoarchitecture.

Exceptions concern one region not previously annotated (*Vsst*, discussed below), and several annotated territories that we found molecularly indistinguishable. For example, *Dld* was similar to more posterior *Dp,* just like *Vd* did not split to *Vs/Vp* in our annotation. While these distinctions may still be relevant at greater resolution or in other contexts, we resorted to a more simplified annotation.

### Unbiased discovery of genes with axial expression

Independent of cell type mapping and regional division, we next aimed to identify genes that carried most weight in spatial segmentation. Spatially restricted genes are of particular interest for species comparisons and understanding developmental origin of cell types. Therefore, we analyzed the topographical distribution of restricted gene expressions according to their medial-lateral, and dorsal-ventral axial extent.

To acquire an unbiased spatial score for each gene, we calculated the spread of lateral-medial (X) and dorsal-ventral (Y) coordinates along the spots that express the gene. Genes encoding high spatial information would localize to a specific territory, and thus show lower-than-expected spread, as compared to dispersed- or sporadic-expressed genes. And just like there may be patterns on the X-Y plane, certain genes are likely to show anterior-posterior patterns (Z), too. This fact may be missed when focusing on a narrower stretch of coronal sections. Therefore, we integrated the X-Y spatial scores for all sections and replicates, revealing an aggregated pattern of top highly axial genes, along the anterior-posterior axis. Axial patterns were not described by a single gene, but several genes shared similar tendencies, which we grouped to modules. We visualized this systematic, integrated data in Fig. 2B.

We found that axially patterned genes showed co-axial correlation; for example, no gene was patterned only dorsal-ventral, but not medial-lateral. Instead, patterned genes always showed axial enrichment on two, or all (X-Y-Z) axes. Further, we observed all possible combinations of co-axial relation. For example, modules 3 and 4 were enriched both anterior and lateral; and either ventral (module 3) or dorsal (module 4). Neuropeptides *PVALB* (module 12) and *CARTPT* (module 2) were localized lateral-dorsal, and medial-ventral; respectively (Fig. 2B-C); and neuropeptide *TAC2* was restricted entirely to anterior-lateral regions.

Overall, our systematic scoring of axial-patterned genes can hint at developmental origins (morphogen-like), and serve researchers when designing new genetic lines.

### GABAergic inhibitory cell types and their spatial distribution in the goldfish telencephalon

As in recently analyzed non-mammalian species ^20–24^, we identify GABAergic cells as those expressing *GAD1*, *GAD2*, and *SLC32A1;* encoding the glutamate decarboxylase enzyme and GABA vesicular transporter, respectively (Fig. 1F). In teleost these genes are known as subpallial markers. In line with this, our spatial transcriptomics data revealed their enrichment in medial, ventral and posterior regions. Within this subpallial, GABAergic class, we observed 40 cell types (Fig. 3A), characterized by specific markers (Fig. 3B-C, and distinct locations in the tissue (Fig. 3D).

**Figure 3:**
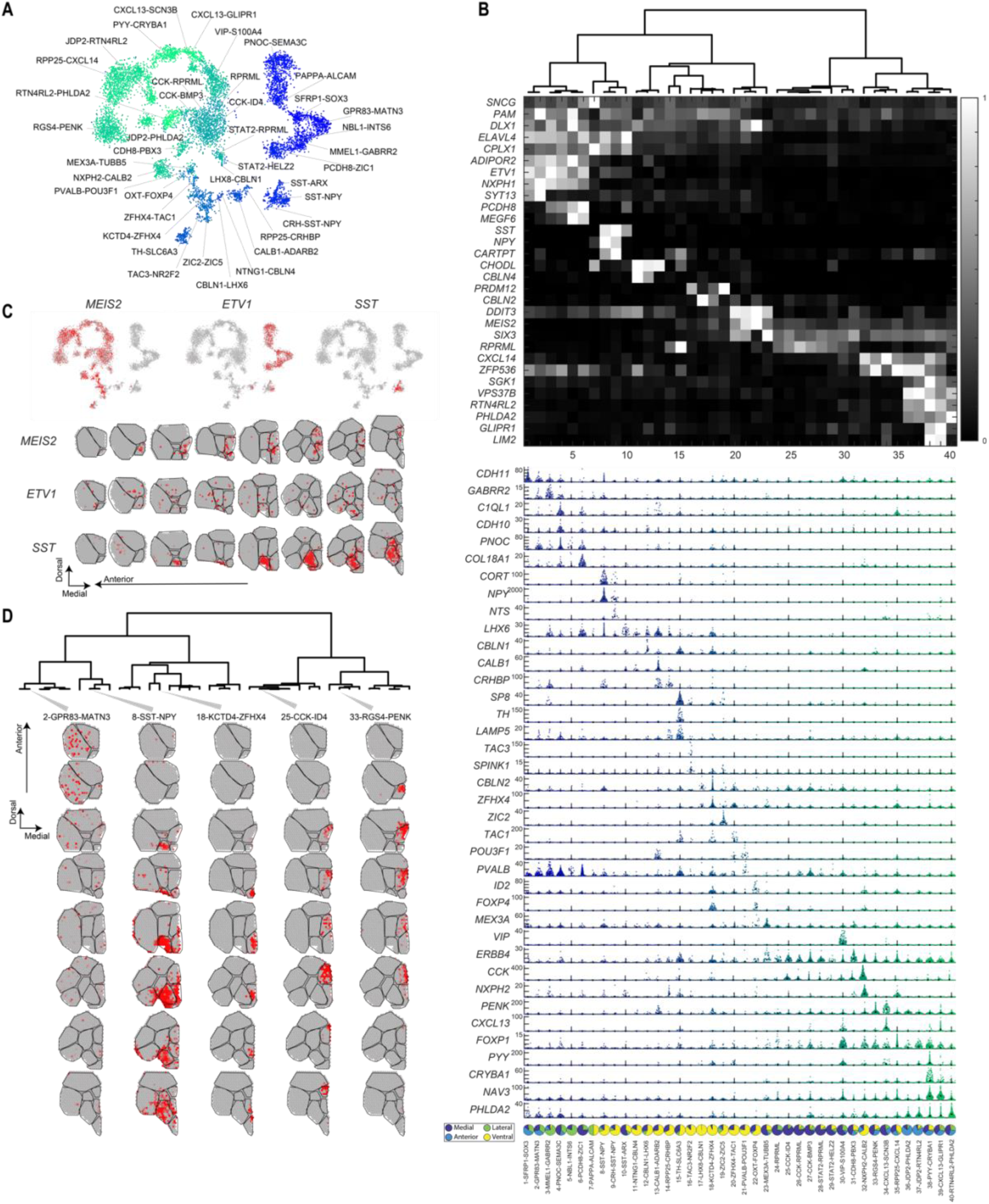
GABAergic neurons in the goldfish telencephalon. **A** t-SNE visualization of GABAergic neurons in the goldfish forebrain. Each dot represents a cell, colored by cell type assignment. **B** All GABA-types arranged in dendrogram order (**GABA1-40**), with top marker gene expression visualized as heatmap (white, high; black, low) and below, violin plots; where each dot represents a single cell; maximum expression (UMI) indicated on the left. Bottom; contribution of four microdissections to each cell type; visualized as pie charts. **C** Expression of three branch-organizing genes, *MEIS2, ETV1* and *SST;* shown in t-SNE as **A** (top, scRNA-seq), and telencephalon coronal hemisphere sections (bottom, spatial transcriptomics); grey, low; red, high. **D** Examples across the GABAergic dendrogram for spatial correlation of Visium spots: five scRNA-seq cell types (columns); across eight a.-p. coronal sections (rows); grey, low; red, high.

First, hierarchical arrangement of the GABAergic cell types on a dendrogram helped reveal their molecular similarities, identify genes that unite related types and gave clues on resemblance to mammals (Fig. 3B-C). For example, cell types on the right-most branch of the dendrogram all expressed mammalian striatal markers *MEIS2* and *PENK*; described below. Further, **GABA1-6** cell types shared expression of transcription factors *DLX1, NXPH1* and *ETV1* and prepronociceptin *PNOC;* all known as cortical interneuron markers in mouse. In mouse, interneurons show a scattered distribution across the neocortex. Mapping goldfish telencephalon **GABA1-6** cell types to the spatial dataset revealed similarly scattered distributions across *Dc* and *Dl* (Fig. 3C-D).

Another example is found in the next branch (**GABA8-10**), which shared expression of *CPLX1* and *ELAVL4* with **GABA1-6.** This related branch also resembled known mammalian inhibitory interneurons by their expression of neuropeptide *SST* (somatostatin). Out of these, **GABA8-9**) additionally expressed a second neuropeptide, *NPY* (neuropeptide Y). Unlike scattered mouse *Sst-*interneurons, however, **GABA8-10** were highly specific to a mid-ventral area in the posterior telencephalon (Fig. 3C-D). We labeled this domain *VSst*, after its most abundant cell type (*SST-NPY*) in our region schema (Fig. 2A). Other prominent markers labelling specific mouse interneuron types (*Pvalb, Vip*), however, behaved entirely different in goldfish telencephalon: They were not specific to interneurons (**GABA1-10**), or even the GABAergic class. For example, the only GABAergic *VIP* population (**GABA30**) expressed no other markers of local interneurons. This cell type was located in the *Vd* compartment (Fig. 3B, Fig. S3); a region we found enriched for neurons resembling an inhibitory *projecting* class (MSN, discussed below). And the classic developmental subpallial divisions of ganglionic eminences-derived interneurons were not apparent in the adult goldfish. Instead, many of the distinguishing markers described in mammals (e.g., MGE, *Sst, Pvalb*; CGE*, Vip, Lamp5, Sncg*) showed ambivalent, overlapping expression.

GABAergic types **GABA11-40** were distributed along the midline; with **GABA11-22** ventrally (*Vv*) and **GABA24-40** dorsally (*Vd*) enriched. Among them, a small population (**GABA15**) resembled mammalian dopaminergic neurons: They specifically expressed marker genes *TH* (tyrosine hydroxylase) and *SLC6A3* (dopamine vesicular transporter); and also *SP8,* a pan-interneuron transcription factor in mouse. Spatially, this cell type appeared in a tiny, ventral-medial nucleus (Fig. 3B, Fig. S3); greatly resembling *th1* expression described in zebrafish telencephalon ^25^.

The final *Vd* clade (**GABA24-40**) was marked by several prominent markers enriched in a projecting GABAergic class termed medium spiny neurons (MSN) in mouse striatum; including transcription factors *SIX3, MEIS2,* neuropeptide *PENK* and dopamine receptor *DRD2* (but not *DRD1*). This raised a possible analogy to the mammalian striatum; but compared to mouse MSNs, these goldfish types had greater diversity. Finally, we identified putative GABAergic neuroblasts along the periventricular zone at the midline, **GABA23**. As described for adult neuroblast in fish ^26^ and the axolotl ^23^, this population specifically expressed *MEX3A* and *TUBB5*.

### Glutamatergic cell types and their spatial distribution in the goldfish telencephalon

We found 48 molecularly defined glutamatergic excitatory cell types (Fig. 4A-B), with distinct, pronounced spatial distributions (Fig. 4C, Fig. S3). They were defined by expression of the glutamate vesicular transporter *SLC17A6* (VGLUT2). More than half of these types additionally expressed *SLC17A7* (VGLUT1) (Fig. 1E). This is in striking contrast to mouse; where *Slc17a6* is reserved almost entirely to the diencephalon, *Slc17a7* dominates the forebrain, and only a minority of cell types co-express the two transporters (e.g., cortex L5, retrosplenial cortex, some amygdaloid nuclei ^27, 28^). In goldfish, *SLC17A6+SLC17A7+* clusters were present in all territories with glutamatergic expression, but they were sparse in *aDl* and *Dld.* Interestingly however, all goldfish telencephalon glutamatergic cells were also marked by *TBR1;* a transcription factor that in the adult mouse is exclusive to cortical pyramidal neurons. Glutamatergic cell types were abundant in pallial territories of the dorsal and lateral telencephalon (e.g., *Dm, Dc, Dl*), with an almost complementary spatial pattern to GABAergic cells, that were highly abundant in subpallial regions (e.g., *Vv, Vd*) (Fig. S3, 4).

**Figure 4:**
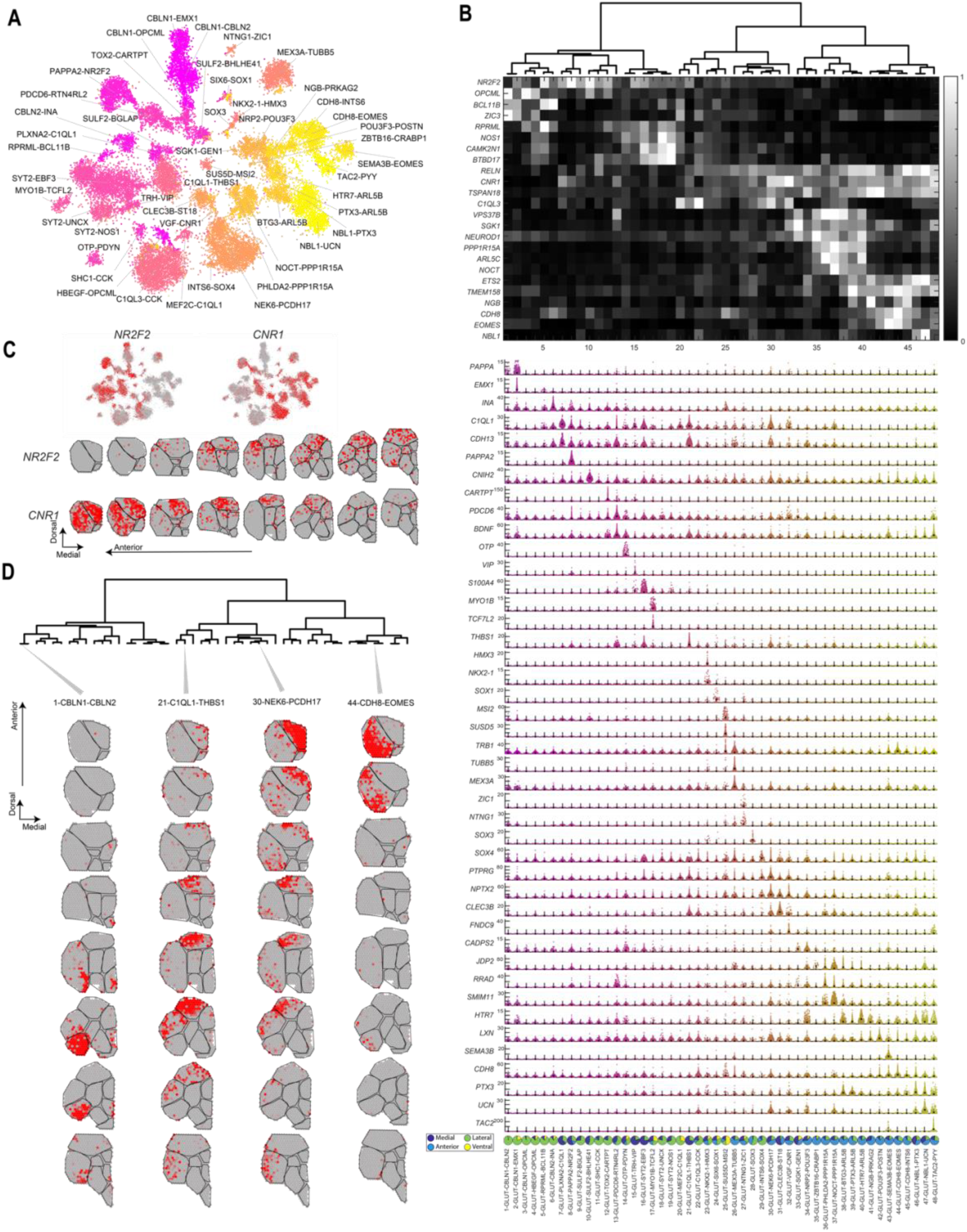
Glutamatergic neurons in the goldfish telencephalon. **A** t-SNE visualization of Glutamatergic neurons in the goldfish forebrain. Each dot represents a cell, colored by cell type. **B** All glutamatergic types, in dendrogram order (**GLUT1-48**), with top marker gene expression visualized as heatmap (white, high; black, low) and below, violin plots; where each dot represents a single cell; maximum expression (UMI) indicated on the left. Bottom; contribution of four microdissections to each cluster; visualized as pie charts. **C** Expression of two branch-organizing genes, *NR2F2* and *CNR1,* visualized on t-SNE, as **A** (top, scRNA-seq) and telencephalon coronal hemisphere sections (bottom, spatial transcriptomics); grey, low; red, high. **D** Examples across the glutamatergic dendrogram for spatial correlation of Visium spots: four scRNA-seq cell types (columns); across eight a.-p. coronal sections (rows); grey, low; red, high.

The first hierarchical division in glutamatergic neurons was reflected in gene expression, and spatial distribution: Here, reelin (*RELN),* cannabinoid receptor 1 (*CNR1),* and thyroid hormone nuclear receptor (*NR2F2)* defined branches of related cell types, with high specificity to the right (GLUT20-48; *RELN, CNR1*) and left (GLUT1-19; *NR2F2*) side of the dendrogram, respectively (Fig. 4B-C). These genes also showed distinct spatial patterns, with little mutual overlap: *CNR1* was anterior, and in *Dm* regions, while *NR2F2* was more posterior, in *Dl* and *Dm* (Fig. 4C). *NR2F2-*types **GLUT1-19** were subdivided into five distinct groups (Fig. 4B). For example, **GLUT1-6** expressed *BCL11B* and populated the ventral *Dl* (Fig. 4B-D and Fig. S3). **GLUT7-9** had a spatial pattern located in the dorsal part of the goldfish telencephalon *Dm* and *Dc* (Fig. S3). The second branch of the dendrogram (*CNR1* **GLUT20-48**) divided into six distinct groups (Fig. 4B). For example, **GLUT29-32** expressed *NEUROD1* and *NOCT,* and were found in the anterior *Dm*; and **GLUT41-48** expressed *EOMES* and were located to the anterior *Dl* (Fig. 4B-D and Fig. S3).

**GLUT14-15** were likely sampled from adjacent hypothalamic regions: These cells expressed the transcription factor *OTP* and the tripeptide hormone *TRH* – genes that faithfully mark several excitatory peptidergic neurons in the mouse hypothalamus ^28, 29^. In the goldfish, we detected these cells in the ventralmost portion of the posteriormost sections (Fig. S3). Finally, we observed a population of glutamatergic neuroblasts: **GLUT26** expressed *MEX3A* and *TUBB5,* like the GABAergic neuroblasts (**GABA23**) did. Unlike **GABA23**, which was restricted to the midline, the glutamatergic neuroblasts lined all pial surfaces (Fig. S3). This pattern strikingly resembled the distribution of goldfish astrocytes (e.g., *HES5+, and MFGE8+, see* Fig. S1). Astrocyte-related *Hes5+* radial glia act as adult neurogenic stem cells in the mouse ^30^. Together, these cell types therefore likely constitute the neurogenic niche on the pial surfaces of the adult goldfish telencephalon ^31, 32^.

### Molecular signatures of teleost telencephalon parcellation

Our analysis of axially patterned genes (Fig. 2), and molecular cell type mapping (Fig. 3-4) allowed to molecularly define regional boundaries. To ease future functional exploration, we next asked the reverse question: Which genes (Fig. 5A-B), and cell types (Fig. 5C), best defined each compartment? For example, the *SST*-dominated nucleus *Vsst* described above was also defined by somatostatin receptor-binding cortistatin (*CORT*). Transcription factor eomesodermin *EOMES* (also known as TBR2) neatly labelled the anterior *Dl*. Calcium binding *NECAB2*, a modulator of the adenosine 2A and mGluR5 receptors, was concentrated in the *Dlv*; but also dispersed, for instance across the *pDm.* In contrast, white matter-dominated *Dc* was indeed best described by myelin binding protein *MBP* and enriched for oligodendrocyte populations. Together, this analysis pinpointed genes that precisely described the molecular base consistent with known anatomical boundaries.

**Figure 5:**
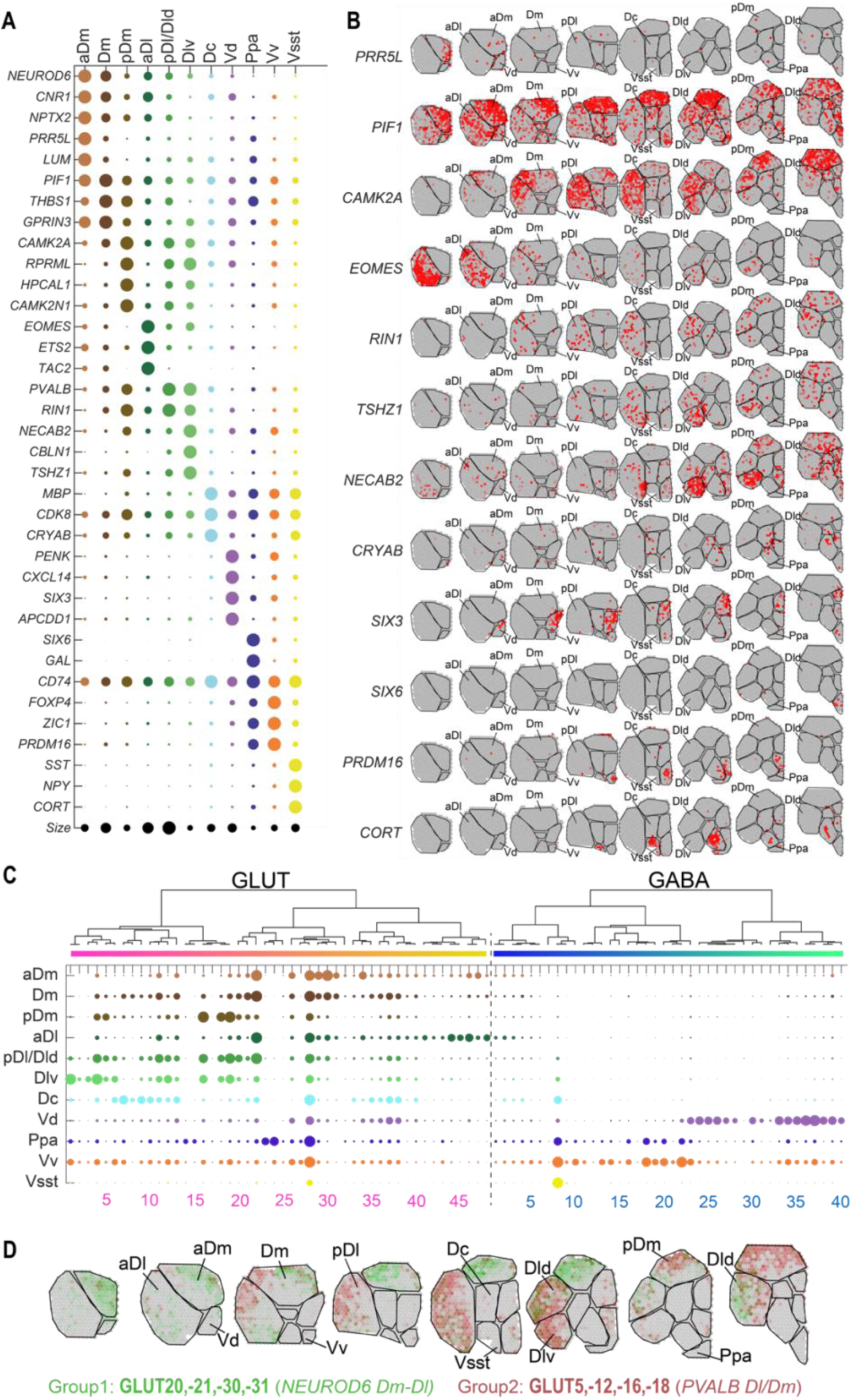
Region-specific marker genes and cell types. **A** Top enriched region-specific marker genes as identified by spatial transcriptomics, presented as dotplot: circle size represents normalized mean expression of each gene in each region. **B** Representative examples of genes from **A**, with spatial distribution of expession across eight a.-p. telencephalon sections; grey, low; red, high. **C** Enrichment of cell types to regions shown as dotplot; where dot size represents the correlation between cell type expression profile and region expression profile. **D** Weighted spatial cell type enrichment for two groups of glutamatergic clusters only (green, **GLUT20, 21, 30, 31**; red, **GLUT5, 12, 16, 18**), indicating a *Dm-Dl* territory switch for both groups, that would result in 3D bends/ horns.

### Excitatory cell types bring a “twist” to axial parcellation

Multiple datapoints in our analysis emphasized the importance of molecular characterization along the full anterior-posterior axis, to better recapitulate underlying structures in 3D. For example, we discovered interesting changes in gene expression and cell type composition between the anterior vs. posterior pallium, in both the *Dm* and *Dl* regions. First, we noticed that *NEUROD6*, a gene known to mark hippocampus CA1 in mammals, was dorso-medial in anterior sections (*aDm, Dm*); but posterior, appeared to “switch” territory to dorso-lateral (*Dld*) (corresponding to Northcutt *Dp* ^18^) (Fig. 2C). Strikingly, neuropeptide *PVALB* appeared to make the opposite “switch”, where more anterior, it was localized dorso-lateral (*Dl/pDl*), but posterior, in the dorso-medial zone (*pDm*) (Fig. 2C). This trend was reflected by additional genes that shared axial patterned expression with *PVALB* and *NEUROD6* (e.g., for *PVALB; CAMK2A, EFNA1, SERPINE1, EMID1*) (Fig. 2B, 5B).

Importantly, spatial distributions of several glutamatergic cell types supported this observation, too: Along the anterior-posterior axis, **GLUT20, -21, -30** and **-31** switched from *Dm* to *Dld,* while **GLUT5, -12, -16**, and -**18**, switched from *Dl* to *pDm* (Fig. 5D, Fig. S3).

In gene expression and cell-type composition – and therefore, likely in functionality – purely axial divisions were thus not always consistent. Instead, we propose that in the current example, the two groups of neuronal populations potentially represent distinct functional units, described by *NEUROD6* (*Dm-Dld*) and *PVALB* (*Dl-pDm*). In three dimensions, each group would form a bend, or horn; analogous to the 3D shape of the rodent hippocampus. Together, these bends would look intertwined, resembling an anterior-posterior “twist” of two molecularly and functionally distinct territories.

### Molecular similarity of cell types between goldfish and mouse indicate conserved brain regions

The structure of the forebrain is diverse across different vertebrates. Today, the mouse brain is of central importance to our understanding of vertebrate nervous systems – but it diverged from teleost about 450 million years ago. Comparative studies on telencephalon of amphibians ^21–23^, bearded dragons ^24^ and turtles ^20^ reveal deeply conserved cell types, even when compared to the mouse brain. Given their vast evolutionary distance, identifying molecular similarities and homologous brain regions between goldfish and mouse may be challenging, but could lead to a better understanding of the evolution of vertebrate brain structure. We therefore continued our investigation of the goldfish telencephalon with systematic comparison to the mouse forebrain. In brief, we integrated our goldfish taxonomy with a published mouse telencephalon cell type taxonomy ^28^. For this purpose, we merged the species’ datasets; to mixed-species datasets of non-neuronal cells, GABA neurons, and GLUT neurons. For each cell class separately, we performed feature selection and dimensionality reduction, and iterated the data integration tool HARMONY ^33^ until embedding of the two species’ cell atlases converged (Fig. S5, Methods). We then applied two parallel approaches aiding comparative analysis: (1) clustering to mixed-species pseudo-cell types, and (2) a *KNN* (k-nearest neighbor) classifier of goldfish cells to mouse cell types (Fig. 6A).

**Figure 6:**
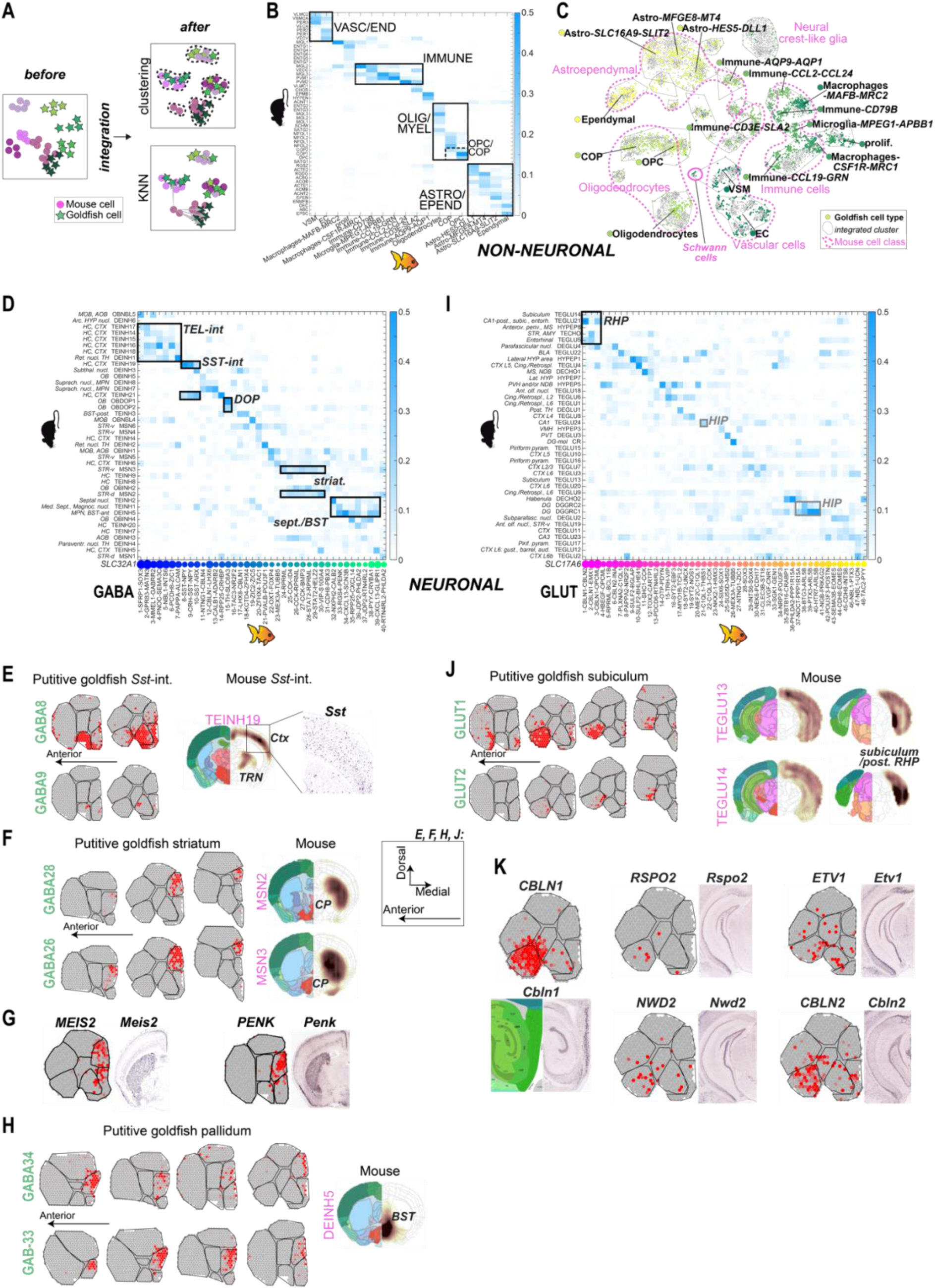
Goldfish and mouse cross species comparison of the telencephalon cell types. **A** Scheme of comparative species analysis between goldfish and mouse^28^ telencephalon. Per cell class, both species’ datasets are integrated (HARMONY), followed by DBSCAN clustering, or a KNN classifier. **B** Non-neuronal cells species comparison visualized as heatmap with conserved cell types outlined (KNN classifier). **C** Non-neuronal cells species comparison visualized as tSNE. Dots represent cells, colored by goldfish cell type; grey, mouse cell; grey outlines, integrated cluster (see Fig. S5); pink outlines and labels, mouse cell class; with mouse Schwann cells highlighted. **D** GABA class species comparison (KNN classifier), with examples for consistency with integrated clusters highlighted. **E** Distribution of *SST-*interneuron types by spatial correlation with goldfish spatial transcriptomics (ST) and mouse ISH^28^; with zoom-in to dispersed *Sst*-expression in isocortex^35^ (Allen Mouse Brain ISH Atlas). **F** Distribution of (putative) striatal neuron types by spatial correlation with goldfish ST (left) and mouse ISH (right)^28^. **G** Expression of two mouse striatum markers in goldfish ST (left) and mouse ISH (right)^28^. **H** Distribution of (putative) pallidal neuron types by spatial correlation with goldfish ST (left) and mouse ISH (right)^28^. **I** GLUT class species comparison (KNN classifier), with examples for consistency with integrated clusters highlighted. **J** Distribution of (putative) retrohippocampal neuron types by spatial correlation with goldfish ST (left) and mouse ISH (right)^28^. **K** Expression of five genes in goldfish ST, and mouse ISH; in (putative) subiculum/retrohippocampal formation.

To circumvent biases inherent to each of the methods (e.g., cluster-size dependency), we present examples of cell types that were in consensus between the two methods, that is, where both approaches identified mouse-goldfish similarities (Fig. 6, Fig. S5). Then, we examined the spatial mapping of these conserved neuronal cell types, to assess to what extent also spatial confines, or brain regions, may have been conserved.

### Conservation of glial cells – and an expansive goldfish immune niche

Non-neuronal cells showed an overall large degree of conservation between the two species, with some interesting exceptions. Among glial populations, we detected three types of astrocytes, that all closely related to the diverse mouse astrocyte populations (Fig. 6B-C, Fig. S5). They expressed a pallet of known mammalian astrocyte markers, including *MFGE8, GFAP, GJA1* (Fig. S1). In addition, goldfish astrocytes expressed genes not typically detected in mouse, e.g., metallothionein *MT4,* delta-like notch ligand *DLL1,* monocarboxylate transporter *SLC16A9* and secreted glycoprotein *SLIT2* (with axon guidance functions in development) (Fig. S1). And unlike in mammals, all populations were almost entirely restricted to the pial and ventricular surfaces (Fig. S1). The population that expressed mouse radial glia marker *HES5* (Astro-*HES5-DLL1)* indeed related closely to both radial glia populations of the two neurogenic niches of the adult mouse, dentate gyrus and subventricular zone (Fig. 6B). Ependymal cells were very closely related between the species in both gene expression and distribution along the ventricles (Fig. 6B, Fig. S5, Fig. S1). This is expected given their necessarily conserved function in circulating cerebrospinal fluid, and the pallet of gene products necessary for this specialization.

The goldfish oligodendrocyte niche reconstituted the entire lineage as described for the adult mouse brain, from oligodendrocyte progenitors (OPC), to committed oligodendrocytes (COP), and finally a mature, myelinating oligodendrocyte (Fig. 6B, Fig. S5, Fig. S1). This observation cements both a conserved developmental origin, and —like ependymal cells, their conserved specialized function, in myelination. Interestingly however, goldfish oligodendrocyte also expressed genes such as *CLDN19, MPZ* and *SPARC* (Fig. S1), that in the mouse are entirely reserved to the myelinating cells of the peripheral nervous system, the Schwann cells ^28^. This is particularly striking given that the mammalian Schwann cells’ developmental origin is in the neural crest, not neural tube. Due to the small amount of Schwann cells included in the mouse dataset however, this unexpected relationship between mouse Schwann cells and goldfish oligodendrocytes was missed by both approaches in our analysis (Fig. 6B-C, Fig. S5).

In contrast to the mouse, the goldfish telencephalon immune niche – marked by *PTPRC* (CD45) – was remarkably diverse (Fig. S1). Like the mouse brain, we detected several types of macrophages (*APOE*) and microglia (*MPEG1*) (Fig. 6B-C, Fig. S1, Fig. S5). Some immune cells bore similarity to myeloid and lymphoid components of the peripheral immune system not generally detected in the mouse brain; such as granulocytes or monocytes (Immune-*CCL2-CCL24*, Immune-*AQP1-AQP9*), B-cells (Immune-*CD79B*) and T-cells (Immune-*CD3E-SLA2*). Therefore, similarity to the annotated mouse forebrain dataset may be misleading (Fig. 6B-C, Fig. S5). For example, granulocyte-like Immune-*AQP1-AQP9* resembled ependymal (EPMB) and subcommissural (HYPEND) cells in *KNN* analysis, and astrocytes (Bergmann glia) in the clustering approach. The putative T-cell Immune-*CD3E-SLA2* resembled vasculature (VLMC/ABC) and epithelium (choroid plexus). And yet, it is tempting to speculate if this observation points to functional analogies over the evolution of cell types in mouse and goldfish.

### SST interneurons: molecular similarity, spatial dissimilarity

In the GABAergic class, species comparison detected putative conserved cell types, with counterparts in both species (Fig. 6D-H). For example, goldfish *SST*-clusters **GABA8-10** were a particularly striking example of a highly conserved neuron type. Both in integrated cluster- and KNN analysis, **GABA8-10** scored closely to mouse cortex somatostatin interneurons TEINH19 and TEINH21 (Fig. 6d, Fig. S5). Yet, as mentioned above, spatial distributions of this neuron type differed in the two species (Fig. 6E): In the mouse, *Sst* interneurons are distributed in a dispersed pattern all over the telencephalon. In the goldfish, we found mapping of *SST*-types **GABA8-10** concentrated, and dominating, in the lateroventral portion of the subpallium, a region we termed *Vsst* (Fig 2A, Fig. S4). *SST* expression patterns both in spatial transcriptomics sections and fluorescent in-situ hybridization confirmed this observation (Fig. 6E, Fig. S6). Therefore, *SST*-interneurons serve as a strong example for a highly molecularly conserved cell type. However, due to their distinct localization, they may nevertheless represent a functionally different type.

In addition, both KNN and integrated cluster analysis confirmed the overall split between interneurons, and putative projecting (MSN-like) inhibitory types. **GABA1-7** resembled a group of cortical and hippocampal interneurons (TEINH14-18); a mix of CGE- and MGE-derived phenotypes (Fig. 6D). The dopaminergic cell type **GABA15** was highly correlated with dopaminergic populations of the mouse olfactory bulb (Fig. 6D, Fig. S5).

### Striatal and pallidal derivatives in goldfish *Vd*

The other branch of goldfish GABAergic cell types (**GABA16-40**) overall had less similarity to local inhibitory cortical interneuron types (Fig. 6D). Instead, some of the top correlated cell types in the mouse included striatal medium spiny cells and inhibitory neurons of the mouse septum and pallidum. **GABA25-28** in the goldfish for example, consistently grouped with mouse medium spiny neurons (MSN) (Fig. 6F). Spatial transcriptomics mapped these cells medial, to the *Vd* (Fig. 2A, purple; Fig. S3). Also, mouse striatum markers *PENK* and *MEIS2* aggregated exclusively in the goldfish *Vd*, forming a continuous regional entity resembling the situation in the mouse striatum (Fig. 6G). Finally, species comparison consistently aligned **GABA33-39** with mouse septal and pallidal inhibitory cell types, and most prominently with BST-population DEINH5 (Fig. 6D, Fig. S5). These cell types were localized in more anterior aspects of the *Vd* (Fig. 6H). Together, these similarities in molecular composition and spatial distribution suggest goldfish homologues to mouse cerebral nuclei in the *Vd*. Our analysis further suggests their resolution to pallidal- and striatal-like regions in the *Vd*’s most anterior, and posterior aspects (including those traditionally termed *Vs* and *Vp*), respectively.

### Glutamatergic neurons suggest a retrohippocampal formation-homolog in goldfish *Dlv*

In contrast to GABAergic, and certainly non-neuronal types, the excitatory cell class showed less coherence between our two approaches to species comparison; implying lower levels of conservation (Fig. 6I-K).

Nevertheless, we found that **GLUT20-22** and **GLUT37-40** were potentially related to hippocampal neurons: Regardless of the analysis approach, both groups were similar to mouse hippocampus proper (Fig. 6I). Yet, this relationship could not be pinpointed to single mouse cell types. Instead, depending on the approach, their relation alternated between hippocampal subdivisions of the mouse (e.g., TEGLU24 (CA1), TEGLU23 (CA3) and DGGRC1-2 (DG)) (Fig. 6I, Fig. S5, Suppl. Table 1). The similarity between these cell types in goldfish and mouse may be partly explained by mouse hippocampal-enriched genes *C1QL1* and *NEUROD6,* that were highly expressed in e.g., **GLUT21-22**. Both groups (**GLUT20-22** and **GLUT37-40**) were located scattered across the *Dm* and *Dl* regions (Fig. S3). Together, these findings hint at limited molecular conservation, and a less spatially defined hippocampal-like formation in goldfish, compared with the mouse.

Most consistently among glutamatergic neurons, one group of goldfish neurons, **GLUT1-2**, scored closely to subiculum glutamatergic cell type (TEGLU14) (Fig. 6I-J, Fig. S5). The goldfish cell type counterparts mapped to the molecularly distinct lateral-ventral region spots, *Dlv* (Fig. 6J, Fig. S3). The similarity between the species was manifested by their shared expression of *CBLN1, CBLN2, NWD2, ETV1* and *RSPO2* (Fig. 6K). In addition, *KNN* analysis only suggested that entorhinal and posterior hippocampal mouse neuron types (TEGLU5 and -21) were closely related, too (Fig. 6I). In mouse, the entorhinal area and subiculum belong to the retrohippocampal formation. They play a major function in hippocampal-cortical interaction, and spatial navigation; e.g., head direction^34^ . Together, these putative subiculum, or retrohippocampal formation neurons suggest an example of conserved molecular identity, corresponding to a regionally confined area, in both species’ telencephala.

## Discussion

Combining spatial transcriptomics with single-cell RNA-sequencing, we generated a neuronal cell type atlas for the telencephalon of goldfish, a carp-like (cyprinid) teleost fish with outwardly-grown (everted) telencephalon characteristic for ray-finned fish (*Actinopterygii*). The resulting datasets will provide a critical resource to analyze neuron type evolution across vertebrates offering to compare huge data sets across various emerging forebrain cell type atlases of mammals ^28, 36^, reptiles ^20, 24^ and amphibians ^21–23^.

Our study discovered 88 GABAergic and glutamatergic neuron types alongside accurate spatial mapping. We determined that all major classes of forebrain cell types found in other species also exist in the goldfish. And in a systematic comparison to mouse telencephalon, many neuronal populations had direct counterparts in the mouse telencephalon – some of which point to conserved brain regions.

Spatial transcriptomics revealed the expression patterns of thousands of genes. We introduced an efficient new way for spatial scoring of axial-patterned gene expression in the telencephalon. This method can be helpful to systematically identify spatially-informative genes. In our case, it identified top potential markers in the mediolateral and dorsoventral axes. Many of these pattern-forming genes were enriched on all three axes – and were thereby in line with anatomical parcellation to molecularly distinct regions, or nuclei. Spatially enriched genes may give clues to underlying developmental processes and evolutionary origins of brain structures in vertebrates.

Beyond spatial enrichment of single genes, we also used *in situ* spatial transcriptomics to map all cell types to their location. Assuming that a molecularly defined neuronal type represents a particular functional unit, we then parcellated the combined map of all cell types, to propose a regional neuroanatomy of the goldfish telencephalon. And while this approach was often in agreement with traditional cytoarchitecture-based telencephalon atlases ^18, 19^, our parcellation revealed a molecular-neuronal, and therefore likely functional base for regional divisions. This speculation was cemented by several conserved cell type homologues revealed in our systematic comparison of goldfish cell types to a mouse telencephalon taxonomy. Here, some conserved neuron types within the goldfish telencephalon showed similar discrete spatial distributions as their counterparts in the mouse forebrain.

For example, brain regions in goldfish putatively homologous striatal domains in mice were identified via cell type similarity between mouse medium spiny neurons and goldfish GABAergic cell types (**GABA26-28**). These populations, and the striatal marker *PENK* specifically mapped to a restricted region of the goldfish dorsal pallium; anterior-posterior *Vd*.

Further, a homolog to the mouse retrohippocampal formation, and in particular the subiculum, was located in a ventral portion of the dorsolateral zone (*Dlv*) of the goldfish. This identification was based on molecular relation between the mouse retrohippocampal cell types together with goldfish cell types **GLUT1-3**. Lesion studies implied *Dlv* in spatial cognition ^8^ previously. And intriguingly, head direction-selective cells specifically, were found both in the rodent subiculum^34^ and in the goldfish *Dlv* ^13^. This reflects an interesting correspondence between the molecular and functional level of these regions.

On the other hand, homology to other parts of the hippocampal formation was less discrete. *NEUROD6*, a clear marker for hippocampus CA1 in mouse, mapped to a putative horn-like shaped *Dm-Dld* region. And in species comparison, we revealed similarities of mouse hippocampus proper neurons to several goldfish cell types, that appeared scattered across the *Dl* and *Dm* regions. This is reflected by contradicting observations in electric fish ^37^ and goldfish ^13–15^, where lesion studies and recordings of space-encoding cells linked all –*Dl, Dm* and even more central telencephalon– to spatial cognition. Each was proposed as a teleost hippocampus homologue. We speculate that together with these functional controversies, our molecular findings point towards a less regionally-bound representation of spatial behaviors than in mammals. Our results contribute a molecular axis to this body of knowledge that may aid future efforts in functional characterization. We anticipate that a multimodular approach, such as the combined use of genetic strategies with functional recordings, and connectivity mapping, will bring decisive insights on conservation.

The integrated analysis of cell types mapping in their spatial context also helped us detect a putative anterior-posterior neuroanatomical “twist”: Moving from anterior to posterior, two groups of molecularly related cell types transitioned from *Dm-Dld*, and *Dl-pDm*. This would have been missed based on pure cytoarchitectural methods, and emphasizes a need for more refined spatial mapping. Additional validation for this structural observation would ideally require a more seamless, full-volume method, able to multiplex enough genes to detect multiple relevant cell types. Since each of the cell type groups are also distinctly marked by the expression of *NEUROD6* (*Dm-Dld*) and *PVALB* (*Dl-pDm*), this could aid their initial functional exploration. For example, it is tempting to speculate whether the “twisted” structure of these compartments may have contributed to the controversy on the location of a functional goldfish counterpart to the hippocampal formation, as discussed above. We suggest that new evidence needs to take into account the precise anterior-posterior alignment, and ideally, combine functional measurements with genetic markers; such as *NEUROD6* (a hippocampus CA1 marker in mouse) and *PVALB* in the above example.

In surprising contrast to the mouse, populations of neuropeptide somatostatin (*SST)*- expressing GABAergic cells formed an entirely homogenous ventral nucleus, that we annotated *Vsst*. *Sst*-neurons in the mouse forebrain function as inhibitory interneurons to cortical, and hippocampal pyramidal cells. But, the nucleus-like aggregation of their goldfish counterparts greatly challenges parallels to their mouse function as local inhibitory cells. Therefore, the *Vsst,* and other *SST*-cells could be of great interest for further exploration; to uncover to what extent this distinct cell type was evolutionary conserved in function and behavior.

The mapping of cell types between goldfish and mouse allowed us to spot possible molecular homologies between brain regions of the two species. Despite their vast evolutionary distance, the plethora of information available about mouse neuroanatomy and cell types enabled us to put the findings in goldfish in a wider neuroethological context. This effort allowed us to identify critical functional components that were preserved along brain evolution. It is important to note that this information does of course not fully describe homologies: We do not follow the developmental nor evolutionary path of each region, and crucial knowledge of regional connectivity needs to be generated in a separate effort. And yet, our approach does not merely test for expression of isolated, selected marker genes; but molecularly defined cell types; that represent functional components of the goldfish telencephalon.

Collectively, our findings provide a comprehensive analysis of the cellular profile and transcriptional architecture of the teleostean telencephalon to reveal neuro-evolutionary similarity with mammalian vertebrate forebrains. We believe that the combined neuroanatomical map, with molecular cell type information, will be of great use for deeper characterization of the regions’ functions. For example, specific populations may be targeted genetically and/or spatially, for recordings during behavioral tasks. In this context, the goldfish is of particular interest, not least due to its large size. To this end, we also established an accessible web resource for free exploration of gene expression across cell types, and spatial context. The data, and this resource could be of value for the goldfish, zebrafish and teleost research community, and open other layers in vertebrate comparative neuroscience.

## Data availability

The sequencing data generated in the current study will be available in the ArrayExpress database at EMBL-EBI before publication. For convenience, the final single-cell expression dataset with annotations and metadata is available as a table at https://storage.googleapis.com/www_zeisellab/goldfish/data_txt_files/goldfish_scRNAseq_data.zip, and our spatial transciptomics (Visium) dataset at https://storage.googleapis.com/www_zeisellab/goldfish/data_txt_files/vis_export_data.zip We provide an online accessible resource for gene- and cell type exploration, for both scRNA-seq, and Visium ST data. The website is temporarily available at https://storage.googleapis.com/www_zeisellab/goldfish/webpilot/main_page_19_2_2023.html#

## Acknowledgments

We gratefully acknowledge financial support from the following funders: A.Z. is supported by the European Research Council (TYPEWIRE-852786), Human Frontiers Science Program (CDA-0039/2019-C) and Israel Science Foundation (2028912). R.S. is supported by the Israel Science Foundation-FIRST program (555/19), Human Frontiers Science Program (RGP0016/2019). H.H. is supported by the Swedish Brain Foundation (Hjärnfonden). T.M. is supported by the COBRE CNAP by the Cognitive and Neurobiological Approaches to Plasticity (CNAP), Center of Biomedical Research Excellence (COBRE) of the National Institutes of Health under grant number P20GM113109, and funds from the Johnson Cancer Center from Kansas State University and HFSP (RGP0016/2019). We thank Dana Sagi and Lior Appelbaum for critical discussion of the data and help with HCR protocol.

## Author contributions

Study design: A.Z. and R.S. designed the study and M.T., S.B., H.H. planned experiments.

Data collection: H.H., S.B., S.G. performed tissue collection and cell preparations, Z.L. performed Visium ST, O.O. performed scRNA-seq and Visium ST, M.T. performed histology.

Data analysis: M.T., S.B., A.Z. wrote all code and analyzed scRNA-seq and Visium ST data.

Data interpretation: M.T., S.B., T.M. H.H., T.S., R.S. and A.Z. critically discussed and interpreted scRNA-seq and spatial spatial transcriptomics data.

Writing: M.T., S.B. drafted the paper, H.H. wrote the paper with help from T.M., R.S., A.Z., and input from all authors.

## Methods

### Animals use

We used adult goldfish (13-15 cm), housed under standard conditions in the aquarium. All experimental procedures followed the legislation under the Israel Ministry of Health Animal Experiments Council and were approved by the institutional Animal Experiments Ethics Committees at the Technion Israel Institute of Technology and Ben Gurion University of the Negev.

### Tissue collection for scRNA-seq and Visium ST

All the experiments were approved by the Ben-Gurion University of the Negev Institutional Animal Care and Use Committee and were in accordance with government regulations of the State of Israel. Goldfish were sacrificed using an overdose of MS-222, followed by transcardial perfusion with freshly prepared, ice-cold, carboxygenated NMDG-based artificial cerebrospinal fluid (aCSF: 93mM NMDG, 2.5mM KCl, 1.2mM NaH2PO4, 30mM NaHCO3, 20mM HEPES, 25mM D-glucose, 5mM Na-ascorbate, 2mM thiourea, 3mM Na-pyruvate, 10mM MgSO4, 0.5mM CaCl2, adjusted to pH7.3-7.4 with concentrated HCl) ^38^. Brains were quickly removed, maintained on ice in aCSF, and superficial meninges and vasculature removed.

### Dissection, cell dissociation and scRNA-seq

For dissociation, the aCSF-perfused brains were quickly embedded in 1.5% Low Melting Temperature agarose, cooled, mounted on a Leica VT1200S Automated Vibrating Microtome and sectioned to 500μm coronal slices (Fib. 1B). Sections were then quickly microdissected in cold aCSF. Microdissected tissue pieces were digested in 800-1000μl papain digest solution (Worthington Papain system, vial 2 reconstituted in 5ml aCSF, and 5% Dnase (vial 3 reconstituted in 500μl aCSF)), 20-25min at 34°C, until mechanical trituration with a wide-diameter fire-polished glass pipette easily separated most of the tissue. Next, remaining undigested pieces were removed by filtering the suspension through an aCSF-equilibrated 30m cell strainer (Partec CellTrix), to a BSA-coated microcentrifuge tube. Cells were pelted 200g 5min at 4°C and resuspended in 200μl aCSF with 2.5% DnaseI (Worthington Papain system, vial 3 reconstituted in 500μl aCSF). For myelin and debris removal, the suspension was layered on top of 1ml 5% OptiPrep (Sigma) in aCSF in a BSA-coated microcentrifuge tube, and centrifuged for 6min 150g at 4°C, with slow ramping. The resulting cell pellet was resuspended in a minimal volume of aCSF, and inspected in a Burker counting chamber for intact cell morphologies and high viability and successful debris removal. At all steps, from perfusion to final single cell suspension, tissue or cells were maintained in ice-cold carboxygenated (95% O2, 5% CO2) aCSF – with the exception of papain digest, where the temperature was 34°C.

Single-cell suspensions were diluted to 1,000-1,000 cells/μl, and processed for 10X Chromium NextGEM generation and scRNA-seq. We followed the manufacturer’s instructions, targeting 5,000-6,000 cells per sample. Sequencing libraries were multiplexed and sequenced on Illumina NextSeq or NovaSeq NGS platforms, targeting a depth of >35-40K reads per cell.

### Quantification and Statistical Analysis

#### Gene orthologous assignment

Orthologs was identified from protein sequence using eggnog-mapper with the EggNOG (v.5.0) orthology data ^39^. In addition, BLASTP^40^ with BLOSUM45 matrix was used in order to identify orthologs using protein sequences, the gene with the higher score was selected. The results from those analyses and the annotation of NCBI were compared and combined. Genes that were found as the same orthologs in both methods got the orthologs name, genes that found just in BLABT got the gene name from BLAST, genes with annotations from NCBI received the gene name from the NCBI annotations. Genes with conflict about the orthology, received different genes from each method, received the serial number from NCBI. Genes which no orthologs or annotations have been found were not included during the further analysis. Most of the orthologs genes we found were one to many.

For the cross-species analysis and correlation, we need one to one orthologs in order to use pairwise correlation between expressions of orthologs genes. Combining those paralogs into a single expression profile was done by summing up the expression of the fish paralogs. Sum of the expression averages noise and gives us the option to perform pairwise correlation between expressions of orthologs across species.

#### Pre-processing and filtering

scRNA-seq data were aligned to the goldfish reference genome and transcriptome (NCBI Assembly ASM336829v1), and the mRNA molecules were counted using the 10X Genomics Cell Ranger (version 5.0.1) ^41^. Molecule counts data from all sequencing runs were merged into one single database which included metadata about each cell. This resulted in 60,705 valid cells. The expression of the paralog genes were summed into one gene. This resulted in 21,427 genes. For normalization and noise reduction we used mean-centering, normalization to a common molecule count, standardization (division by the standard deviation) and log transformation.

#### Features selection and dimensionality reduction

With any type of clustering the choice of feature space is crucial. Highly variable genes (features) were selected using coefficient of variance as a function of mean (CV) ^42^. Genes detected in fewer than 20 cells or more than 60% of all cells will be marked invalid and will not be used in the analysis of the highly variable genes selection, in addition we excluded the immediate early genes. Briefly, we calculated log2 of mean and log2 of CV per gene, then we fitted a linear line to the data using linear regression, and selected the genes having the greatest offset from the fitted curve; this would correspond to genes with higher-than-expected variance. The number of the selected genes (cut off) for the analysis was determined using “elbow plot”, a common heuristic to choose a cutoff point. Then, for dimensionality reduction Principal Component Analysis (PCA) was used based on the highly variable genes. The number of the principal components (PCs) used for the downstream analysis was determined using “elbow plot”.

#### Clustering and classification

Clustering of all cell at once suitable for finding major cell types, but not optimal for finding finer subdivisions among cells of the same kind (e.g., <u>interneurons</u> in a dataset containing both neurons, vascular cells and glia). We decided to use a multi-level clustering approach, first split cells by major class and then splitting the major classes into subclass, the clusters were annotated based on known cell type specific markers. In each level the feature space was selected according to the entire set of cells, projected by PCA. In order to exclude batch effects and technical effects from biological ones, we run the Harmony ^33^ on the PCA, an algorithm that projects cells into a shared embedding in which cells are grouped by cell type rather than dataset specific conditions, thus obtaining a new PCA. 2D embedding using t-distributed stochastic neighbor embedding (t-SNE) was performed on the PCA space. Then the cells were clusters based on their distance in 2D using the Density-based spatial clustering of applications with noise (DBSCAN) non-parametric algorithm ^43^.

#### Level 1 analysis

We pooled samples by tissue and performed Pre-processing, clustering, classification, gene enrichment, and marker gene detection (see below for details on these procedures).

#### Level 2 analysis

We split cells by major class and manually annotated clusters to indicate major classes of cells: *Neurons, Glutamatergic Excitatory and GABAergic Inhibitory and Non-Neurons*. We also manually identified and removed clusters that were clearly doublets between these major classes (e.g., *Vascular-Neurons*) as well as clusters that were of poor quality.

#### Level 3 analysis

We split cells from the major classes into subclasses separately using the same analysis steps as for Level 1 and 2. Despite our efforts, there remained still some clusters that were suspected doublets, as well as over-split clusters that lacked clearly defining gene expression differences. To create the final consolidated dataset, we extensively annotated and named each cluster according to enriched genes and known cell type specific markers. The level 3 analysis was the basis for all downstream analysis.

#### Visium protocol

Goldish are perfused trans-cardially with ice-cold artificial cerebrospinal fluid (aCSF) then OCT frozen brains are sectioned by the cryostat and mounted coronally as an intact tissue on a Visum Spatial slides arrays of capture area, each capture area has thousands of barcoded spots containing millions of capture oligonucleotides with spatial barcodes unique to that spot. Afterwards, we stain the tissue with DAPI staining and image the mounted tissue prior to dissociation using a brightfield microscope and DAPI suited laser using a fluorescent microscope. Then, the tissue is permeabilized to release mRNA from the cells, which binds to the spatially barcoded oligonucleotides present on the spots. A reverse transcription reaction produces cDNA from the captured mRNA. The barcoded cDNA is then pooled for downstream processing sequencing.

#### Spatial transcriptomics (ST) Visium (10x Genomics)

For spatial transcriptomics, freshly perfused brains were quickly coated,- and cryomold-embedded in cryoprotective OCT (TissueTek), and flash-frozen in isopentane equilibrated on dry ice. We maintained brains in sealed bags at -80C until processing. Per brain, we collected eight left hemisphere coronal cryosections at 10μm thickness, aiming at approximately 200-300μm spacing, spanning the anterior-posterior axis of the telencephalon, onto the Visium ST gene expression slide (4 capture areas, 10x Genomics). For tissue preparation, we followed the manufacturer’s instructions, with the following specifications: methanol fixation, immunofluorescence staining with DAPI only (no antibody), imaging at 10x magnification on a Nikon Eclipse Ti2: DIC, DAPI, and TRITC channels for fiducial and section alignment and 25min permeabilization, as we had previously determined on test sections, using the Tissue Optimization kit. We then proceeded with the Visium Gene Expression Kit following the manufacturer’s instructions, with 15 PCR cycles for cDNA amplification. Sequencing was performed on Illumina NGS platforms to a depth of 150-200M reads per sample (i.e., capture area, or section).

#### Spatial analysis

With the exception of the spatial barcodes instead of cell barcodes (spots instead of cells), most of the steps discussed in the next paragraph are similar to scRNA-seq analysis pipeline and discussed in detail in the single cell analysis section.

Briefly, after pre-processing, filtering, normalizing and feature selection, we perform dimensionality reduction by principal component analysis (PCA) for the high dimensional data that we get from the sequencing output, followed by 2D embedding with t-distributed stochastic neighbor embedding (t-SNE) based on their similarity in the high dimensional gene expression space. This resulted in a 2D map. Focusing on neuronal markers, we clustered these spots according to their distance in 2D using the density-based spatial clustering of applications with noise (DBSCAN) algorithm. Thus, spots with similar gene expression are grouped together in the same cluster.

We also use K Nearest Neighbors (KNN) to regroup the outlier spots from the DBSCAN algorithm into their closest neighbor cluster. Afterwards, these spots could be remapped to their original location on the tissue based on the spatial barcode index, to test whether clusters had an unbiased spatial distribution in the tissue.

#### scRNA-seq integration to spatial transcriptomics

Spatial transcriptomics (ST) spots contain RNA transcripts from multiple cells, rather than single cells, hence we need to deconvolute this mixture of cells in order to compare it with scRNA-seq annotated cell type data.

We then conduct a feature selection of highly variable genes for scRNA-seq and ST dataset alone, then we combine and intersect the two feature selection gene lists together. After normalizing both datasets, we conduct a smoothed combinatorial scoring of the expression of the top 10 genes in each cell type into the cluster of each pair of single cell types and Visium spatial spots. The output result gives us the rho-the correlation results for each spot with each cell type.

We then map those scores in the original positional location in the actual X-Y position of the section. We also create a weighted colormap for the sections, (based on the same colormap of the single cell types) using the weighted score of all cell types in all spots. In order to see a global pattern comparison, we normalize the rho correlation values and present all the section spots together in a heatmap form.

#### Axial spatial genes scoring

The spatial scoring values are based on the normalized standard deviation of the gene expression of each expressed gene in the goldfish telencephalon from the centroid mean X Y Z axes of the Visium section spots; we discovered a clustered pattern of selected top highly axial genes. Spatial axial score for each gene was calculated a follows for mediolateral (sigma x) and dorsoventral (sigma y):

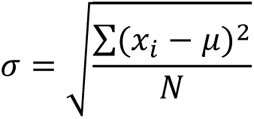

σ = specific required gene standard deviation.

N = number of spots with required gene expression in each Visium section.

Xi = each expression value of the required gene in the Visium section.

Μ = axial centroid mean of X or Y axes for all spots in the individual Visium section.

We also calculated the combined overall axial spatial score for both mediolateral and dorsoventral axes scores together:

σoverall = √ (σx^2 + σy^2)

#### Cross species comparison and data integration

After annotating cell types, we integrated our goldfish taxonomy with our published mouse telencephalon cell type taxonomy ^28^, using the data integration tool HARMONY ^33^, separately for each main cell classes; GABA neurons, GLUT neurons, and non-neuronal cells. First, we concatenated the two species’ datasets, normalized them, and removed genes that were not intersected between the two species. Then, we selected highly variable genes from each species individually, and combined the two gene lists for feature selection. Next, we normalized the merged datasets and performed dimensionality reduction in PCA space, to exclude batch effect and technical effects from biological ones. Then, we performed HARMONY in PCA space. This projected cells into a shared embedding in which cells are grouped by cell type rather than dataset-specific conditions, and resulted in a new PCA that accounts for experimental factors, but not biological ones.

We then proceeded with two methods to aid in the comparative analysis of goldfish and mouse cell types: DBSCAN clustering and KNN (k-nearest neighbors).

In the KNN approach, goldfish cells were classified for their k-nearest mouse cell neighbors in the PCA space, where we calculated the fraction of each specific cell type according to their shared mouse cell type that they were classified to, All KNN classifiers presented used k =25 (where k is the number of mouse cell neighbors), although testing different k-values (10, 50, 100) did not majorly impact classification fractions, that remained stable.

In the clustering approach, the new PCA space was then embedded in t-SNE, and clustered using DBSCAN algorithm. Similar cell types of both species fell in close proximity to their counterparts, in the Euclidean 2D space embedding. Finally, we calculated the percentage of each cell type in each of the integrated clusters and sorted the cell type according to their percentage values. When a mouse cell type has high similarity or analogy to a goldfish cell type, we expect a high percentage for both in an integrated cluster.

