## Supplemental Figures for "A telencephalon cell type atlas for goldfish reveals diversity in the evolution of spatial structure and cell types"

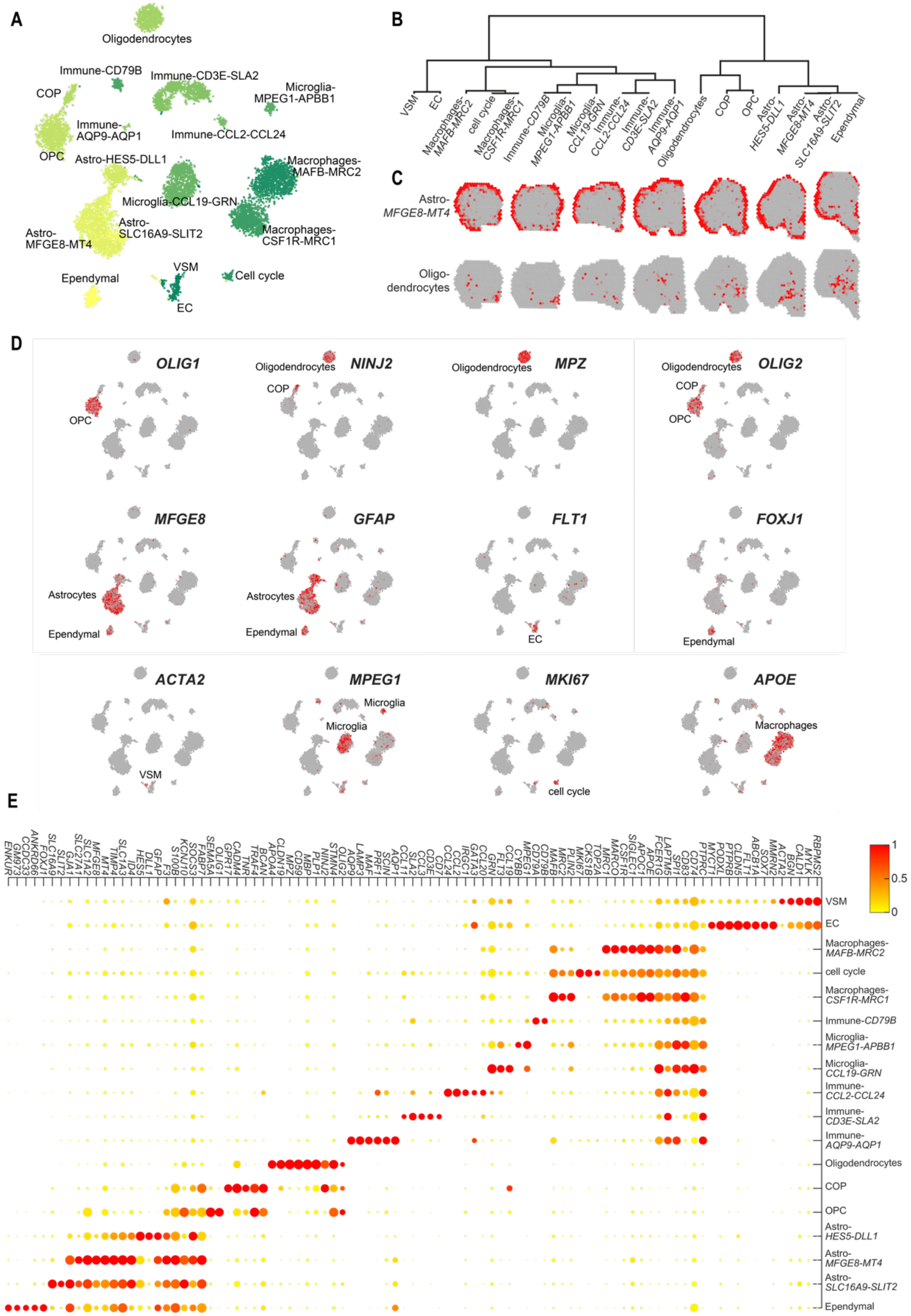


**Figure S1: Non-neuronal cells in the goldfish telencephalon. A** t-SNE visualization of non-neuronal cells in the goldfish forebrain. Each dot represents a cell, colored by cell type. **B** Top marker gene expression, visualized on t-SNE plane (as **A**); grey, low; red, high. **C** Top: Dendrogram of all non-neuronal cell types. Bottom: Examples across the non-neuronal dendrogram for spatial correlation of Visium spots: two scRNA-seq cell types (rows); across eight a.-p. coronal sections (columns); grey, low; red, high. **D** DotPlot visualization of top marker gene; where dot size represents the percentage of cells in a cluster expressing each gene; dot color represents the normalized average expression level of each gene.

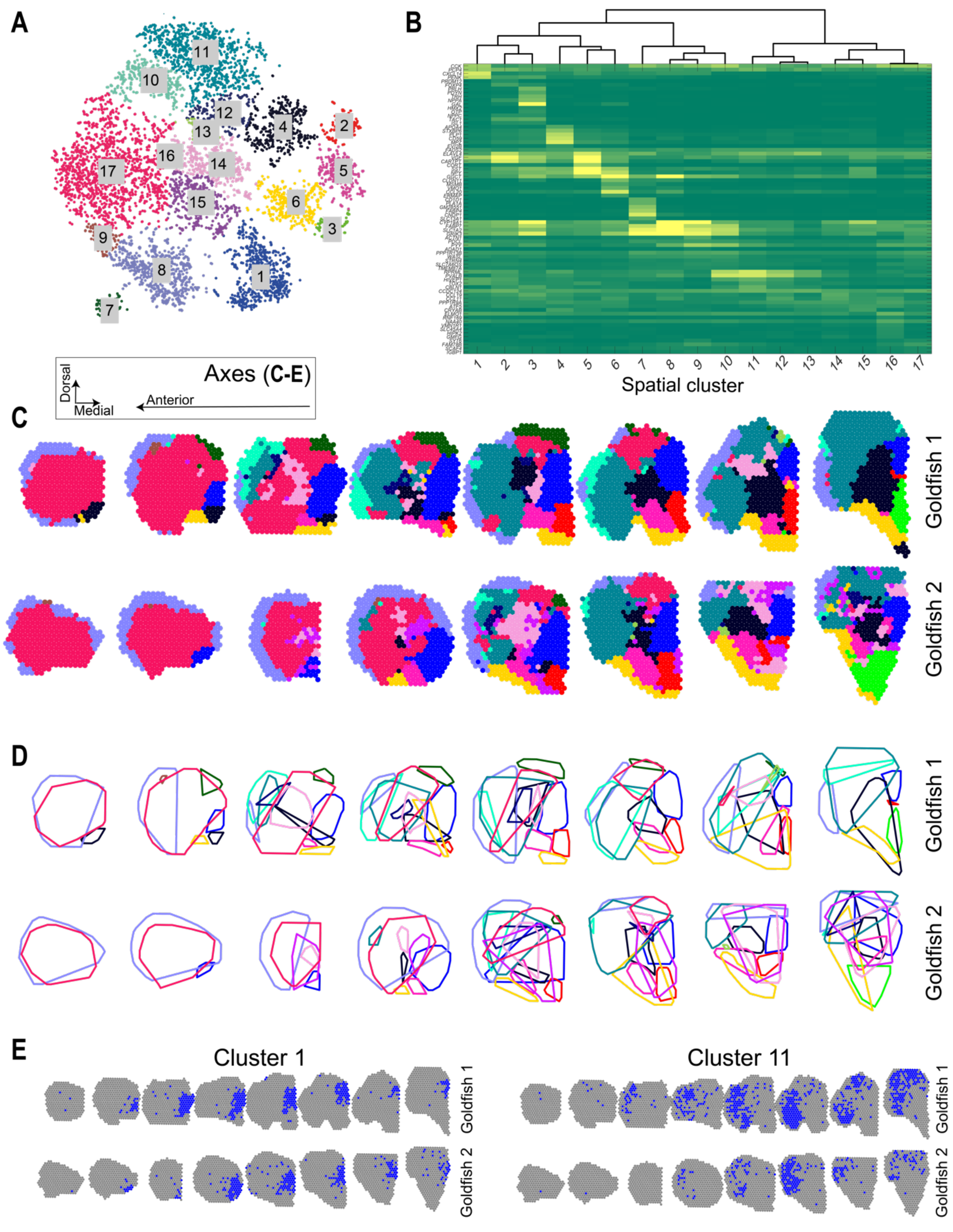


**Figure S2: Spatial transcriptomics reveals distinct regional neuroanatomy across the goldfish telencephalon.** **A** t-SNE visualizing 6,710 Visium spots (=spatial mini-bulk samples) for two goldfish forebrains; each dot represents a spot, colored by spatial cluster. **B** Heatmap of spatial cluster gene expression, for top marker genes (rows) of the 17 spatial clusters (columns). Green, low; yellow, high expression; expression normalized per row. Spatial clusters are arranged by hierarchical clustering; see dendrogram above. **C** Visualization of Visium spots in their spatial sampling location on the sections, colored by spatial cluster assignment (as **A-B**). **D** Visualization of cluster outlines in the sections, as seen in **C**. **E** Visualization of two examples for spatial clusters; cluster 11 (left) and 1 (right); with spots assigned to each cluster shown in blue; grey, all Visium spots.

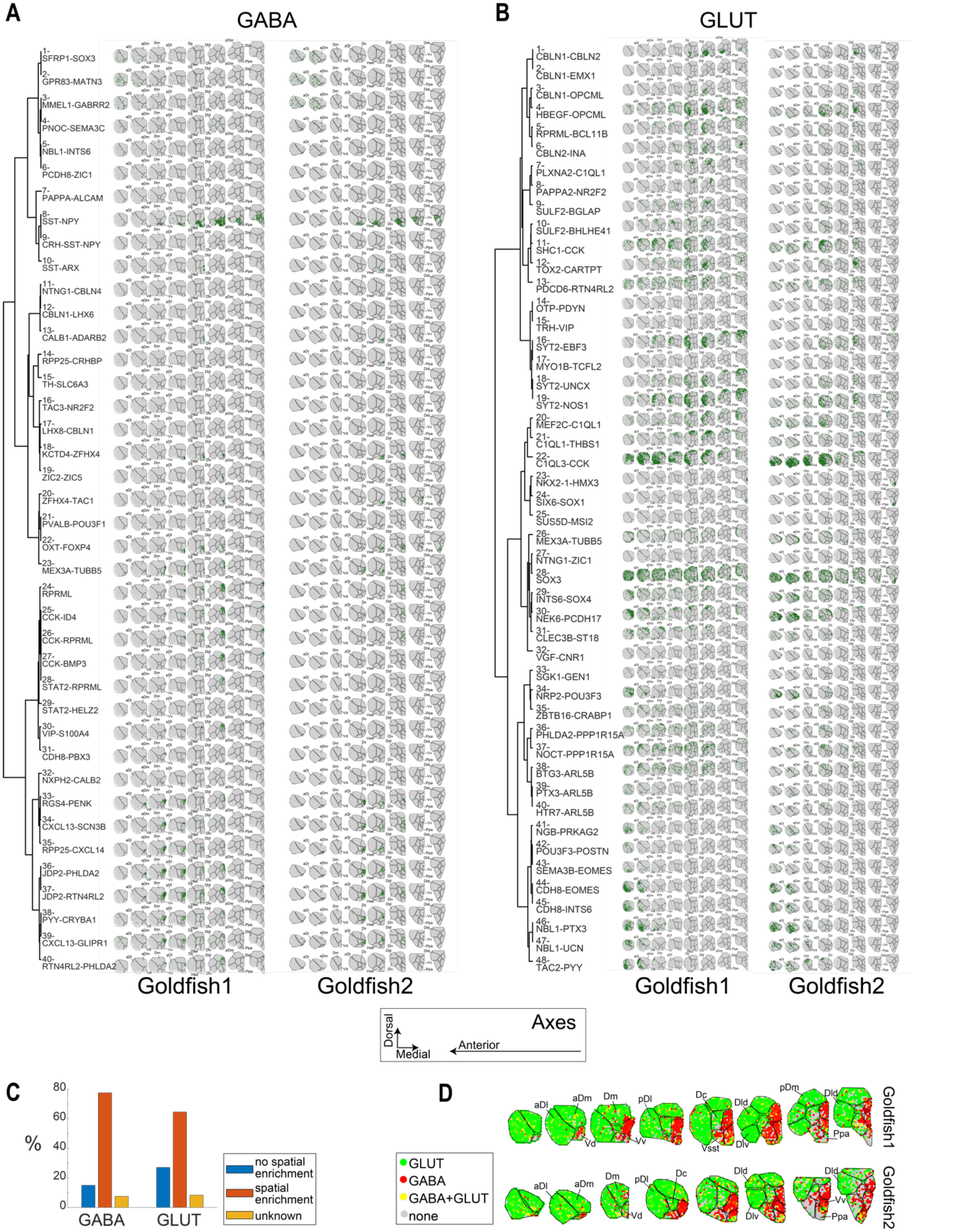


**Figure S3: Cell types spatial mapping to Visium spots in goldfish telencephalon. A** Localization of all GABAergic cell types (rows; in hierarchical cluster order) to Visium spots; by enrichment score overlaid on anterior-posterior sections for goldfish 1 (left) and 2 (right); green, high; grey, low. **B** as **A,** for glutamatergic cell types, in hierarchical order. **C.** Bar plots summarizing spatial enrichment counts for GABA and GLUT cell types. **D.** Glutamatergic (green) and GABAergic (red) expression patterns across the Visium sections spots.

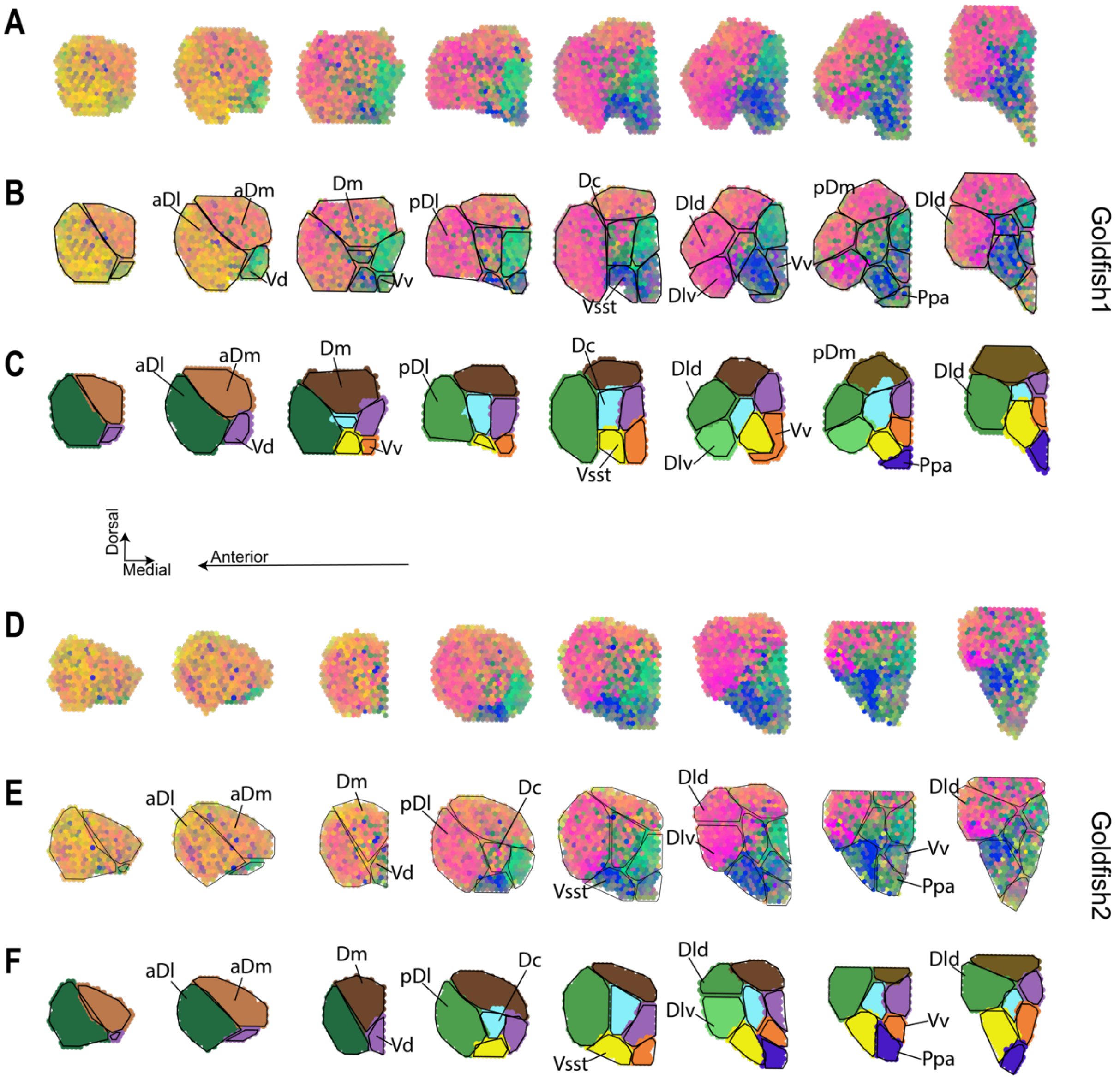


**Figure S4: Molecular neuronal cell types-based neuroanatomical parcellation in two goldfish telencephala. A-C** Goldfish 1; **D-F** Goldfish 2. **A, D** Weighted colormap of goldfish telencephalon cell type mapping to location within Visium spots. **B, C** Weighted colormap like **A, D**; with regional parcellation border and names indicated. **C, F** Regional parcellation schematic, where each color indicates a different region across the AP axis. Suggested names in B-C and E-F are based on similarity to Northcutt 2006 [18] region names. Axes directions shown in inset window.

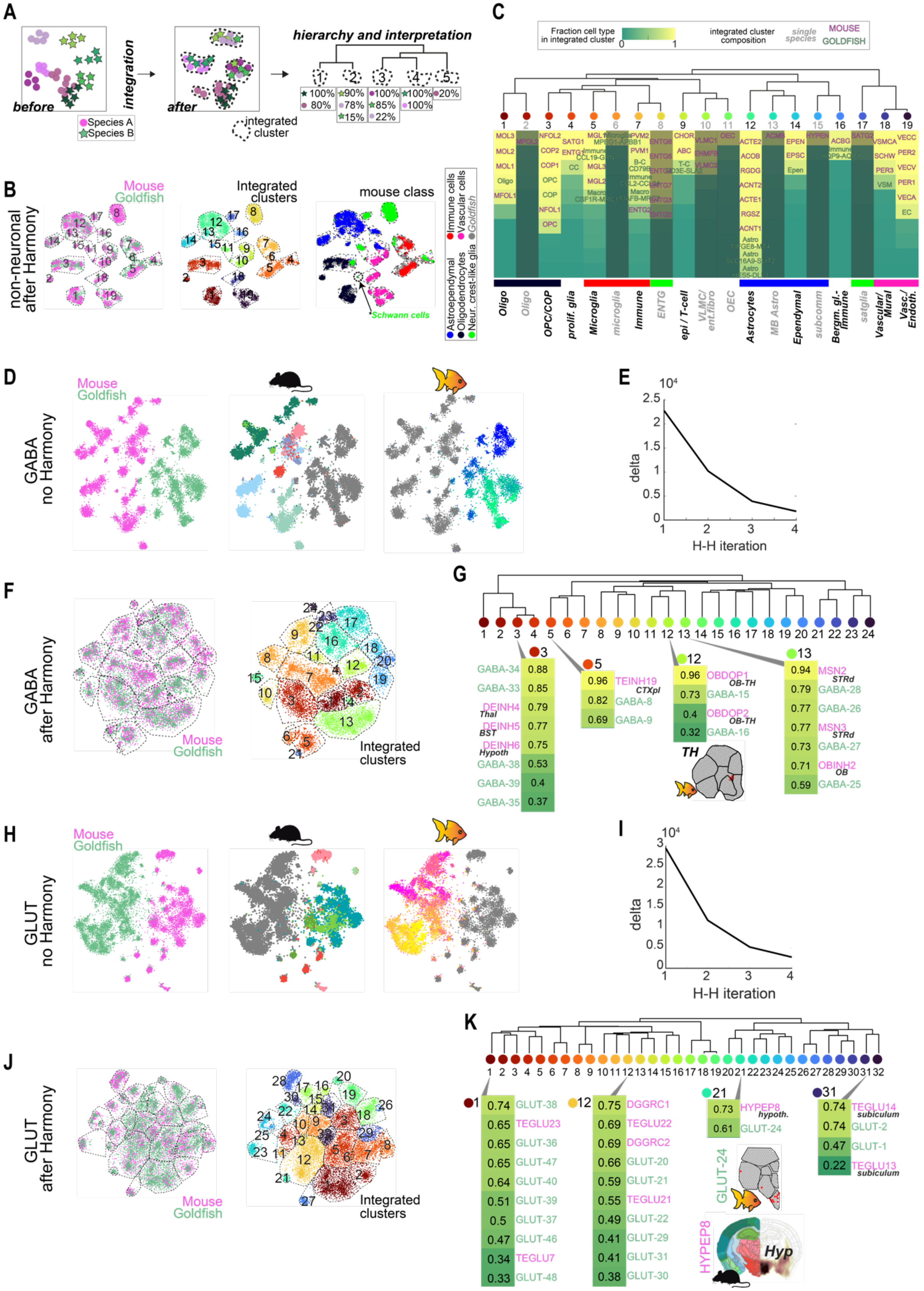


**Figure S5: Species comparison of mouse and goldfish telencephalon cell types. A** Schematic of species comparison analysis as in Fig. 6a, including integrated clustering analysis interpretation. **B** t-SNE visualizations of the combined goldfish-mouse scRNA-seq non-neuronal datasets, after integration using HARMONY. Each dot represents one cell; colored by species (left), integrated cluster assignment (center) and mouse cell class (right). The location of mouse Schwann cells is indicated on the rightmost tSNE. **C** Composition of 19 non-neuronal integrated clusters as in **A**, in hierarchical order (see dendrogram, top). Mouse and goldfish cell types contributing to each integrated cluster are indicated, where the heatmap represents fraction of cells of each annotated cell type contributing to the integrated cluster. Below, interpretation of integrated cluster. Grey, integrated cluster is made up of a single species. Species comparison of GABAergic (**D-G**) and glutamatergic (**H-K**) cell types; **D,H** tSNE visualization before HARMONY, each dot represents one cell, colored by species (left), mouse cell origin by Allen Mouse Brain atlas region color annotation (center), and goldfish cell type (right). **E,G** Mean projection Euclidean distance between each two consecutive iterations of HARMONY, starting with high arbitrary distance value (delta), 5 H (Harmony iterations) in total to plateau. **F,J** tSNE visualization after HARMONY, each dot represents one cell, colored by species (left), and integrated cluster identity (right). **G,K** Dendrogram of integrated cluster analysis like in **C**, with examples of putative conserved cell types highlighted.

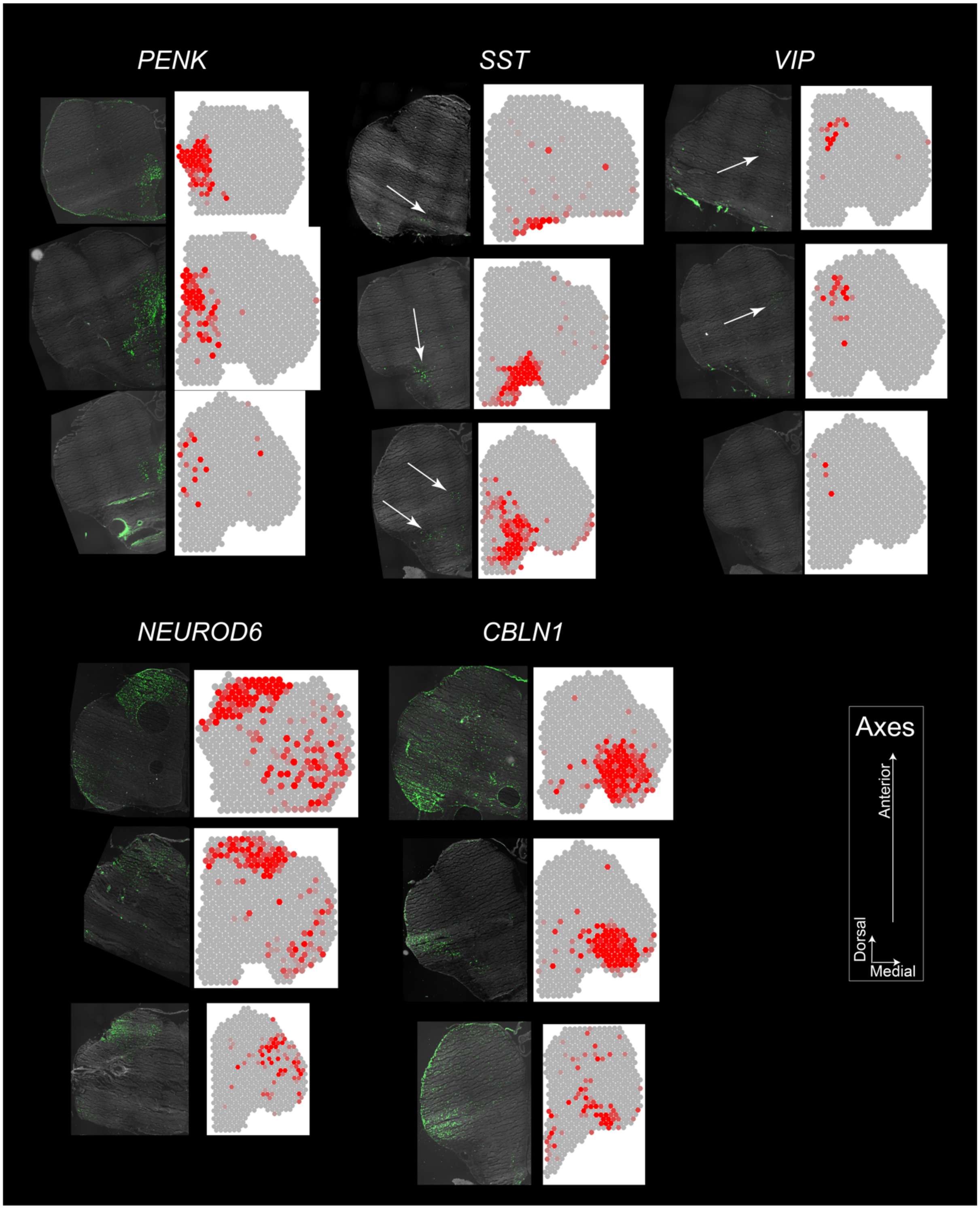


**Figure S6:** Validation of Visium gene expression data (right) with fluorescent *in situ* hybridization by hybridization chain reaction (HCR RNA-FISH, left); for three anterior-posterior sections per gene (*PENK, SST, VIP, NEUROD6, CBLN1* as indicated). Visium and HCR-FISH sections are approximately matched for anterior-posterior position, and shown mirrored to mimic right-left hemispheres. Arrows point out sparse positive signal.
